# Graph analysis of the guilt network highlights associations with subclinical anxiety and self-blame

**DOI:** 10.1101/2024.02.21.581371

**Authors:** Michal Rafal Zareba, Krzysztof Bielski, Victor Costumero, Maya Visser

**Affiliations:** Neuropsychology and Functional Neuroimaging Group, Department of Basic and Clinical Psychology and Psychobiology, Jaume I University, 12-006 Castellon de la Plana, Spain; Institute of Psychology, Jagiellonian University, 33-332 Krakow, Poland

**Keywords:** guilt, anxiety, MRI, structural connectivity, functional connectivity, graph theory

## Abstract

Maladaptive forms of guilt, such as excessive self-blame, are common characteristics of anxiety disorders. The associated network includes the superior anterior temporal lobe (sATL), underlying the conceptual representations of social meaning, and fronto-subcortical areas involved in the affective dimension of guilt. Nevertheless, despite understanding the anatomy of the guilt processing circuitry, network-level changes related to subclinical anxiety and self-blaming behaviour have not been depicted. To fill this gap, we used graph theory analyses on a resting-state functional and diffusion-weighted magnetic resonance imaging dataset of 78 healthy adults. Within the guilt network, we found increased functional contributions (higher clustering coefficient, local efficiency and strength) of the left sATL for individuals with higher self-blaming and trait-anxiety, while functional isolation (lower clustering coefficient and local efficiency) of the left pars opercularis and insula was related to higher trait-anxiety. Trait-anxiety was also linked to the structural network’s global parameters (mean clustering coefficient), with the circuitry’s architecture favouring increased local information processing in individuals with increased anxiety levels. Previous research suggests that aberrant interactions between conceptual (sATL) and affective (fronto-limbic) regions underlie maladaptive guilt and the current results align and expand on this theory by detailing network changes associated with self-blame and trait-anxiety.

## 1. Introduction

During our lives we find ourselves navigating through the complex world of interpersonal interactions on a daily basis. Accurate comprehension of the actions and messages conveyed by others, as well as the expectations and the norms of acceptable behaviour on our side, constitute the key building blocks for everyday functioning. As a result of these interactions, our perceptions of them and the associated mental events, we experience a variety of social feelings. One type of these, labelled ‘moral’, enables us to be motivated by the needs of other individuals or environment in the absence of benefit to ourselves or our kins (as reviewed in Zahn et al., 2020). At the times when we believe we have transgressed the rules of acceptable behaviour or have behaved immorally, we often experience feelings of guilt, which are thought to serve as a punishment cue and facilitate inhibition of such actions in the future (Monteith, 1993). Effective functioning of this system is thus essential for operating in our everyday lives. Not surprisingly, mental health conditions associated with difficulties in social functioning, such as depression and anxiety, are characterised by the prevalence of maladaptive forms of guilt, such as excessive self-blaming (Kim et al., 2011; Candea and Szentagotai-Tătar, 2018). Given that the subclinical populations may share common symptomatology with the diagnosed individuals (Besteher et al., 2017), investigating the neuronal correlates of the feelings of guilt in the former group may provide us with an invaluable insight on the functioning of this circuitry in general and additionally increase our understanding of how these symptoms emerge in the clinical populations.

Owing to the recent meta-analytic and review efforts (Bastuni et al., 2016; Gifuni et al., 2017; Eslinger et al., 2021), it has been established which brain regions participate in processing the feelings of guilt. Some key areas in this network include the frontopolar cortex, the subgenual cingulate cortex and adjacent septal region (SCSR). In addition, Eslinger et al. (2021) draws special attention to the right superior anterior temporal lobe (sATL) given that the connectivity between this region and SCSR during guilt processing is altered in the major depressive disorder (MDD) patients and has the ability to predict illness recurrence (Green et al., 2012; Lythe et al., 2015). With sATL suggested to be involved in accessing the conceptual representations of social meaning, and the activation of fronto-subcortical areas reflecting the affective dimension of guilt, it has been proposed that aberrant interactions between semantic and emotional networks might underlie the prevalence of maladaptive guilt across mood and anxiety disorders (Green et al., 2012; González-García and Visser, 2023).

Nevertheless, despite the general understanding of the anatomy of the guilt processing circuitry, as well as the functions the individual regions may play in the phenomenon, currently it is not known whether and how their network-level interactions are in fact associated with self-blaming behaviour. The works reported above have concentrated on task-related measures of local activity and connectivity between two specific regions rather than looking at the guilt processing areas as a functional and structural brain network. Describing the physiology and anatomy of the brain utilising approaches such as graph theory is especially important given that no single area functions in isolation and the more complex characteristics, apparent only at the network level, might be otherwise easily overlooked in the traditional mass-univariate analyses. To fill in this crucial gap, we used a resting-state functional magnetic resonance imaging (rs-fMRI) and diffusion-weighted MRI (dMRI) dataset to construct the functional and structural guilt networks and investigate the associations of self-blaming emotion regulation strategy with their global and local graph theory parameters. Taking into account the multiple co-occurring mental processes that give rise to blaming oneself, including self-awareness, theory of mind and conflict monitoring (Gifuni et al., 2017), it is plausible that variability in the functional or anatomical connections of brain regions governing any of these functions could shift the delicate balance in this complex psychological machinery and contribute to increased self-blaming tendency. Alternatively, such an observation could also be related to the characteristics of the guilt network as a whole. Indeed, individual differences in other emotion regulation strategies have been associated with both nodal and global graph theory parameters (Pan et al., 2018; Jacob et al., 2019), and thus we hypothesised to observe a similar pattern for the self-blaming behaviour.

Furthermore, despite the importance of maladaptive guilt in anxiety disorders (Candea and Szentagotai-Tătar, 2018), no study so far has examined how the neural correlates of guilt processing vary in regards to individual anxiety levels. To explore this uncharted area of research, we set out to investigate how the global and local graph theory parameters of the functional and structural guilt network would be associated with trait anxiety in a non-clinical population. Anxious people have been reported to have differential structural and functional connectivity both within and across the canonical brain networks (Xu et al., 2019; Yang et al., 2020). The areas implicated in guilt processing map onto several of these networks and as such it would be difficult to predict the direction of the combined impact of such distinctions on both the local and global graph parameters. Taking into account the prevalence of excessive guilt in anxious individuals, we hypothesised that individual differences in trait anxiety would be reflected in both types of network parameters, similar to the results of the previous whole-brain and network-specific studies (Tao et al., 2015; Zhu et al., 2017; Makovac et al., 2018; Guo et al., 2021).

Therefore, the objectives of the current study are to define the guilt network using the available literature, and use graph theory to examine the characteristics of this circuitry that are associated with self-blaming tendency and subclinical anxiety. Based on previous research, we expected the guilt network, and in specific the sATL and medial frontal regions, to interact differently in individuals with high self-blame and anxious personality traits.

## 2. Methods

### 2.1. Dataset

The project was performed using the fully anonymised MRI and behavioural data obtained from the Max Planck Institute Mind-Brain-Body Dataset (Babayan et al., 2019). After applying the initial inclusion criteria, i.e. right-handedness, secondary or higher level of education, no current medication use and no history of neurological, psychiatric and substance use disorders, 95 out of 153 available young participants (aged 20-35) were found eligible for the analyses. Data from 17 subjects were discarded due to excessive head motion, making the final sample of the fMRI analysis equal to 78 (20 females and 58 males). To achieve the correspondence between the functional and structural neuroimaging indices, we analysed the dMRI data only from the previously mentioned 78 individuals. Two subjects (1 female and 1 men, both aged 20-25) were found ineligible due to the lack of fieldmap data and a visible scanning artefact, making 76 the final sample of the dMRI investigation.

Trait-anxiety was assessed using the State-Trait Anxiety Inventory (STAI; Spielberg, 1989). The scale is made out of 20 Likert-type items scored from 1 (almost never) to 4 (nearly always), making the outcome range from 20 to 80. In turn, the self-blaming emotion regulation strategy was tested using a four item subscale of the Cognitive Emotion Regulation Questionnaire (CERQ; Garnefski et al., 2001). Each of these items was scored on a 5-point Likert scale from 0 (almost never) to 4 (almost always), making the possible range of scores from 0 to 16. In the case of both tools, the higher the individual score, the more pronounced the psychological feature. STAI and CERQ scores were moderately correlated in our sample (ρ = 0.403; p < 0.001). The full characteristics of the cohort is provided in Table 1.

**Table 1.** Demographic summary of the sample. The presence of ^#^ indicates non-normal distribution of the data and the sole use of median and range.

| Variable | Mean/median (SD) | Range |
| --- | --- | --- |
| Age | 20-25, N = 42<br>25-30, N = 31<br>30-35, N = 5 | 20-35 |
| STAI | 37.22 (8.04) | 21-56 |
| CERQ self-blaming subscale | 4 (n/a) # | 0-11 |

Abbreviations: SD, standard deviation; n/a, non-applicable; STAI, State-Trait Anxiety Inventory; CERQ, Cognitive Emotion Regulation Questionnaire.

### 2.2. MRI data acquisition

Three types of neuroimages acquired with a 3 Tesla scanner (MAGNETOM Verio, Siemens, Healthcare GmbH, Erlangen, Germany) equipped with a 32-channel head coil were used in the course of the investigation. High-resolution anatomical data was collected with T1 MP2RAGE sequence (176 sagittal slices; 1 mm isotropic voxel size; TR = 5000 ms; TE = 2.92 ms; flip angle 1/flip angle 2 = 4°/5°; GRAPPA acceleration factor 3). The T2*-weighted resting-state fMRI data was acquired with a gradient echo planar multiband imaging sequence (64 axial slices acquired in interleaved order; 2.3 mm isotropic voxel size; TR = 1400 ms; TE = 30 ms; flip angle = 69°; multiband acceleration factor = 4). During the resting-state scan, the participants were instructed to lie awake with their eyes concentrated on a low-contrast fixation cross. 657 volumes were collected for each individual, resulting in a scan duration of 15 mins 30 s. Additionally, the high angular resolution diffusion-weighted images (HARDI) of the whole brain were collected axially (88 slices in interleaved order) with a 1.7 mm isotropic resolution (60 diffusion-encoding gradient directions, b-value of 1000 s/mm^2^, 7 b0 images, TR = 7000 ms, TE = 80 ms, FA = 90°, GRAPPA acceleration factor = 2, 32 reference lines, multiband acceleration factor 2).

### 2.3. MRI data preprocessing

The preprocessing of T1-weighted data was performed in the Computational Anatomy Toolbox (CAT12; Gaser et al., 2022). Individual anatomical images were skullstripped and segmented into grey matter (GM), white matter (WM) and cerebrospinal fluid (CSF). The skullstripped T1-weighted data was used later for the delineation of regions of interest (ROIs) in each participant, as well as the coregistration with the other MRI modalities.

The preprocessing stream of the functional data consisted of steps available in the FSL (susceptibility distortion correction; Jenkinson et al., 2012), R (motion scrubbing using fMRIscrub package; Power et al., 2012) and AFNI (all other steps; Cox, 1996). The first five volumes of the fMRI time series were discarded to allow for signal equilibration. Then the data underwent despiking, and slice-timing, motion and susceptibility distortion corrections. At this point the functional neuroimages were coregistered with the skullstripped anatomical data. Subsequently, the fMRI data was prewhitened, detrended, denoised and band-pass filtered in the 0.009–0.08 Hz range. The regressors in the denoising matrix included: six motion parameters and their derivatives, as well as WM and CSF signal. Additionally, volumes with extensive motion (i.e. framewise displacement > 0.25 mm) were censored. Participants with a mean framewise displacement higher than 0.2 mm or more than 30% of the censored volumes were excluded from the further analysis. The minimal length of the non-censored time series was thus 10 min 51 s, which is sufficient for a reliable estimation of the functional connectivity (Van Dijk et al., 2010). In order to ensure the spatial independence of the signal in all the nodes of the network no smoothing was applied.

The dMRI data was preprocessed using MRtrix3 software (Tournier et al., 2019). First, the images underwent denoising and unringing, which was followed by distortion, eddy currents and motion correction using the dwifslpreproc wrap-up, which makes use of the dedicated functionalities available in FSL (Jenkinson et al., 2012). Subsequently, the diffusion images were bias field-corrected. The response functions for WM, GM and CSF were estimated with the use of the dhollander algorithm (Dhollander et al., 2019), and they served as an input for the single shell 3 tissue constrained spherical deconvolution (SS3T-CSD) performed using MRtrix3Tissue (https://3Tissue.github.io), a fork of MRtrix3 (Tournier et al. 2019). Following intensity normalisation, anatomically constrained tractography (ACT; Smith et al., 2012) was performed based on the coregistered 5 tissue image generated by the *5ttgen fsl* function. Compared to the standard fibre tracking algorithms, ACT is a much more biologically reliable method, given that the connections are seeded from the GM-WM interface and terminated upon either entering cortical GM or within subcortical GM, which is in agreement with the characteristics of neuronal projections derived from histological studies. In each participant, 10 million streamlines were generated using the iFOD2 (improved 2nd order integration over fiber orientation distributions) algorithm. The default settings were applied with the exception of the maximal track length reduced to 100 mm (compared to the default 170 mm, given the data resolution) and the cutoff value set to 0.06. As the last step, the generated streamlines were refined with the SIFT2 algorithm (Smith et al., 2015) to allow for the use of the number of streamlines as a valid index of the structural connections density.

### 2.4. Guilt network construction

The regions (nodes) of the guilt network were selected based on a recent meta-analysis of studies investigating neural correlates of guilt (Gifuni et al., 2017) and a review on the functional neuroanatomy of moral social feelings (Eslinger et al., 2021). We identified 19 ROIs, which were primarily planned to be centred on the reported peak coordinates. Note that for the peak coordinates of the Eslinger et al. (2021) review, we reported the values mentioned in the original studies (Zahn et al., 2009; Moll et al., 2006; Green et al., 2009). Given the fact that the peaks of five areas mapped onto WM and in the case of the fMRI analysis the majority of the signal of interest is believed to originate in GM, their coordinates were slightly adjusted to ensure the proper GM coverage. ROIs were primarily delineated in the MNI space as 3.45 mm radius spheres (1.5 times the fMRI data voxel size; see Table 2 for their mass centre coordinates). To refine the functional ROIs to include only GM voxels, we: 1) averaged the CAT12-derived normalised GM segments across all the participants; 2) excluded voxels with values lower than the threshold used in the volumetric anatomical analyses (i.e. 0.2); and 3) binarised the resulting image and used it to as an inclusive mask for the previously defined ROIs.

**Table 2.** The nodes of the guilt network.

| Location | MNI coordinates | Reference |
| --- | --- | --- |
| R medial orbitofrontal cortex * | 11, 51, -3 | Eslinger et al., 2021<br>Zahn et al., 2009 |
| R subgenual cingulate cortex<br>R septal area | 4, 15, -5 | Eslinger et al., 2021<br>Moll et al., 2006 |
| L subgenual cingulate cortex<br>L septal area | -4, 15, -5 | Eslinger et al., 2021<br>Moll et al., 2006 |
| L anterior cingulate cortex | -4, 28, 24 | Gifuni et al., 2017 |
| L superior frontal gyrus | -4, 56, 22 | Gifuni et al., 2017 |
| L medial frontal gyrus (BA 6) | -4, -6, 48 | Gifuni et al., 2017 |
| L medial frontal gyrus (BA 10) * | -12, 46, 8 | Gifuni et al., 2017 |
| L medial frontal gyrus (BA 9) | -4, 42, 40 | Gifuni et al., 2017 |
| L insula *<br>L inferior frontal gyrus pars opercularis * | -46, 8, 5 | Gifuni et al., 2017 |
| R inferior frontal gyrus pars opercularis | 46, 12, 6 | Gifuni et al., 2017 |
| L superior temporal gyrus | -46, -60, 20 | Gifuni et al., 2017 |
| L middle temporal gyrus | -62, -12, -10 | Gifuni et al., 2017 |
| L superior anterior temporal lobe | -50, 4, -10 | Gifuni et al., 2017 |
| R superior anterior temporal lobe | 57, -3, -6 | Eslinger et al., 2021<br>Green et al., 2009 |
| L posterior cingulate cortex *,#<br>L parieto-occipital sulcus *,# | -15, -56, 7 | Gifuni et al., 2017 |
| L precuneus | -4, -54, 38 | Gifuni et al., 2017 |
| L supramarginal gyrus * | -50, -38, 27 | Gifuni et al., 2017 |
| L cuneus | -4, -92, 10 | Gifuni et al., 2017 |
| R lingual gyrus | 4, -78, 2 | Gifuni et al., 2017 |

Abbreviations: L, left; R, right; BA, Brodmann Area.
\* The area originally mapped onto white matter and thus had its location slightly adjusted.

Delineation of ROIs in the individual brains was achieved by warping them from the MNI space. The spatial correspondence of the network nodes with the fMRI and dMRI data was achieved using the matrices created during the linear coregistration of T1 images with the respective image of other modality.

In the case of the functional network, the nodes’ time-series were extracted and the functional connectivity matrix was calculated using Pearson correlation. Negative correlations were removed from the resulting matrices to ensure easier interpretability of the topological measures (Qian et al, 2018). To normalise the distribution of the correlation values, Fisher z transformation was applied. Such values served as the edges of the functional network. In the case of distance-based graph measures, the obtained z-scores were inverted (1/z) to ensure their proper calculation (Rubinov and Sporns, 2010).

For the structural connectivity data, the ROIs were dilated by 1 voxel to ensure that they covered the border of GM and WM, given that in ACT this interface serves as the seed for tractography (Smith et al., 2012). In each subject, the resulting nodes were inspected visually, and if required, corrected manually to ensure that they were restricted to the areas of interest. The structural connectivity matrix was created by summing up the SIFT2-generated (Smith et al., 2015) weights of all the streamlines connecting every pair of nodes. The edges of the structural network were defined in the same manner as for the functional data.

Weighted, undirected graphs were created using the matrices derived from the fMRI and dMRI data in R (version 4.3.0) using the igraph package (Csardi and Nepusz, 2006). As the graph measures are sensitive to network density (Yeh et al., 2021), the analyses were repeated using sparsity ranging from 5 to 30% with 5% steps.

### 2.5. Graph theory measures and statistical analysis

The same graph parameters were calculated for both the functional and structural networks. On the nodal level strength, weighted clustering coefficient, betweenness centrality and local efficiency were extracted. As for the global network parameters, modularity (Leiden algorithm; Traag et al., 2019), mean clustering coefficient and global efficiency were calculated. The definitions of these measures are provided in the Supplementary Table 1 (Barrat et al., 2004; Freeman, 1979; Latora and Marchiori, 2001; Traag et al., 2019).

Pearson or Spearman correlations, depending on data distribution, were calculated between the graph indices and psychometric measures, separately for STAI and CERQ scores. The significance was assessed with non-parametric permutation testing (5000 permutations) using the jmuOutlier package (Garren, 2017) in R (version 4.3.0). The nodal results were Bonferroni-corrected for multiple comparisons (family wise error rate; FWE) at the level of p_FWE_ < 0.05.

Apart from testing for the associations of the graph measures with the psychometric scores, the betweenness centrality of the nodes was used to examine whether the areas pinpointed by the aforementioned analyses functioned as hubs of the guilt network. These results were, however, treated as secondary and they were thus included in the Supplementary Material (Tables 2-3 and Figures 3-14).

## 3. Results

### 3.1. Functional network

For the functional network, significant associations between the questionnaire scores and graph measures were observed for two nodes: the left sATL (Figure 1); and the left inferior frontal gyrus pars opercularis (IFGpo) and insula (Figure 2). At 25% sparsity the local efficiency (ρ = 0.37; p_FWE_ = 0.027) and clustering coefficient (ρ = 0.357; p_FWE_ = 0.027) of the left sATL were both positively correlated with the strength of the self-blaming coping strategy. The association between the local efficiency of the left sATL node and CERQ score was further replicated at 30% sparsity (ρ = 0.349; p_FWE_ = 0.034).

**Figure 1.**
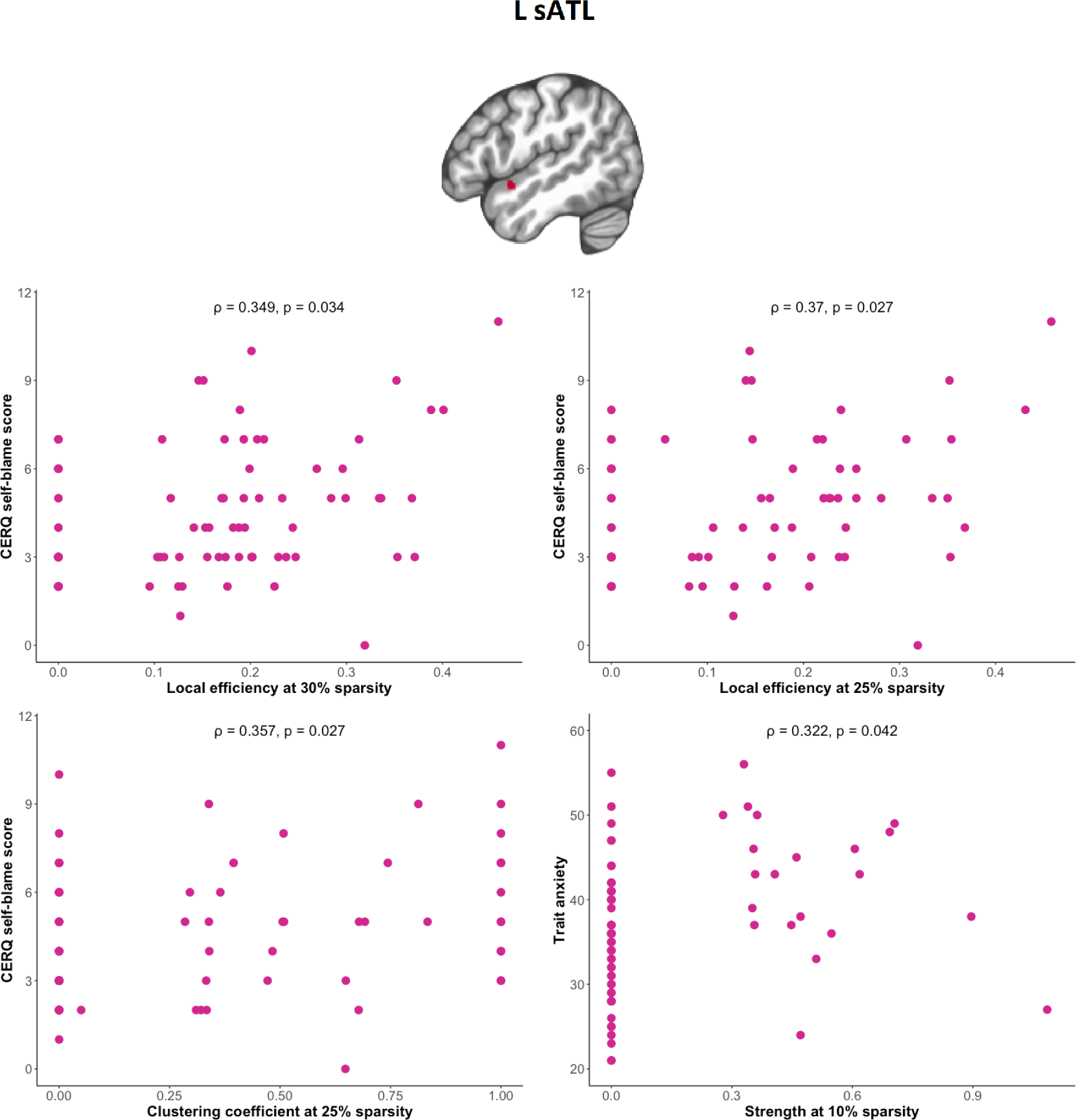
The associations of the functional network-derived graph metrics of the left superior anterior temporal lobe (L sATL) with trait anxiety and self-blaming tendency as measured with Cognitive Emotion Regulation Questionnaire (CERQ; Garnefski et al., 2001).

**Figure 2.**
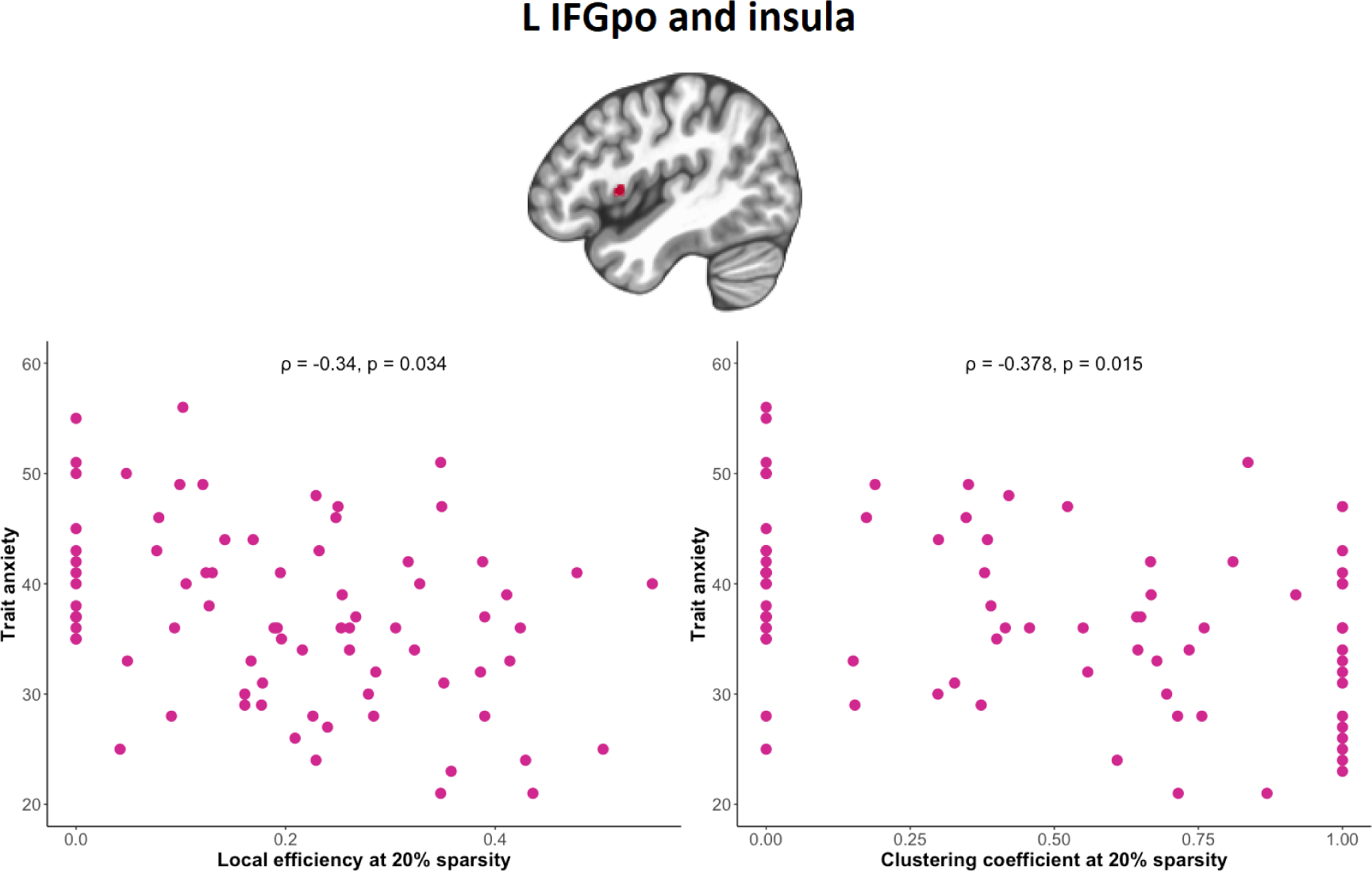
The associations between trait anxiety and the functional network-derived graph properties of the left inferior frontal gyrus pars opercularis (L IFGpo) and adjacent insula.

**Figure 3.**
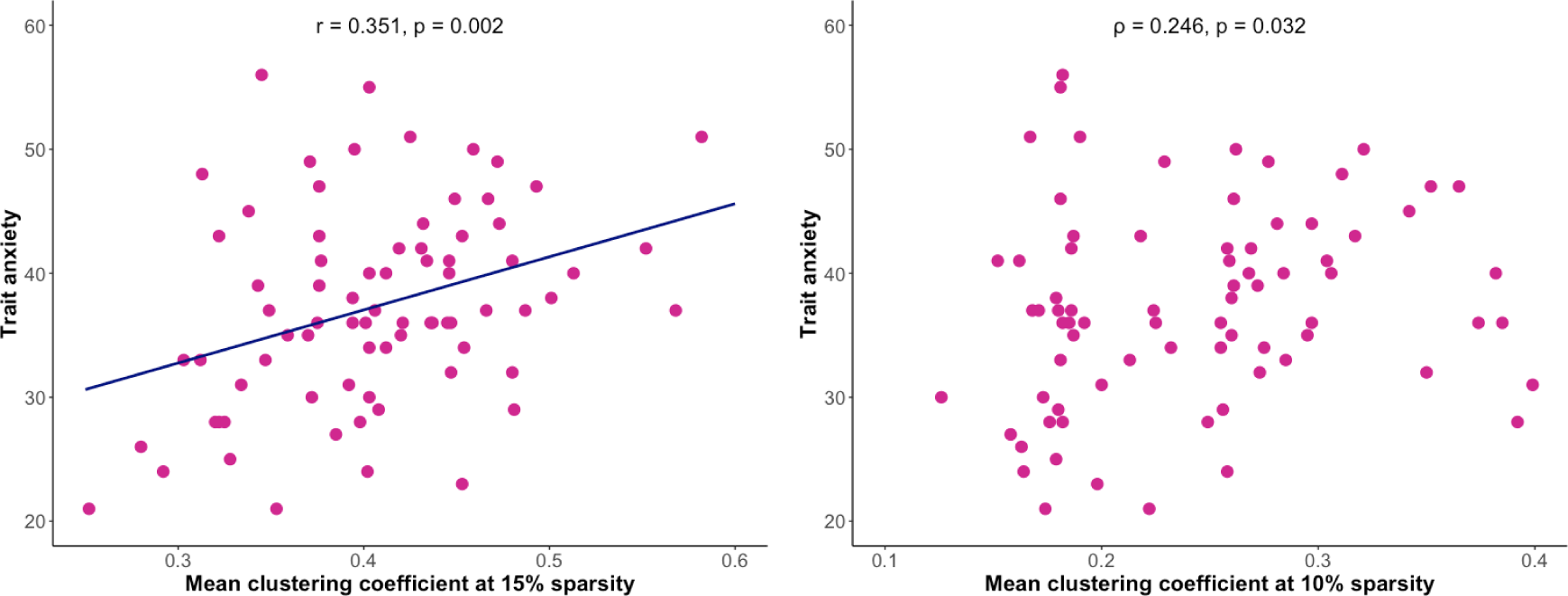
The associations between the mean clustering coefficient of the structural guilt network and trait anxiety.

Interestingly, at 10% sparsity the strength of the same region was positively correlated with trait anxiety (ρ = 0.322; p_FWE_ = 0.042). Additional associations with the STAI scores were observed at 20% sparsity for the left IFGpo and insula, namely the local efficiency (ρ = −0.34; p_FWE_ = 0.034) and clustering coefficient (ρ = −0.378; p_FWE_ = 0.015) of the node were both negatively correlated with trait anxiety. Neither the left sATL, nor the left IFGpo and insula were found to function as hubs of the functional network at the aforementioned sparsities (Supplementary Table 2, Supplementary Figures 4 and 6-8). Associations between the global graph properties and psychometrics were not found.

### 3.2. Structural network

In the structural analyses, associations were found between global metrics and anxiety, and nodal parameters and self-blaming tendency. Mean clustering coefficient of the network was positively associated with anxiety at 10% (ρ = 0.246; p = 0.032) and 15% density (r = 0.351, p = 0.002; Figure 3). Additionally, at 30% sparsity the clustering coefficient of the left medial frontal gyrus (Brodmann area 10) was positively related to self-blaming behaviour (r = 0.338, p_FWE_ = 0.042).

Aligning with the methods used on the functional network, the influence of head motion was controlled. Maximal displacement across all the diffusion-weighted volumes was calculated for each participant, and correlated with the highlighted significant results. The only metric confounded by motion was the clustering coefficient of the left medial frontal gyrus (Brodmann area 10) at 30% sparsity (ρ = 0.23, p = 0.046). The lack of such associations for the global network results (p > 0.05) confirms their validity.

## 4. Discussion

The functional network analyses revealed that self-blaming behaviour and anxiety are more pronounced in individuals with more strongly interconnected left sATL, while higher levels of trait anxiety are further associated with increased functional isolation of the left IFGpo and insula. Moreover, anxiety is positively linked to the structural network’s mean clustering coefficient, indicative of a neural architecture favouring increased local information processing. These results thus partially confirm the hypotheses, linking self-blaming emotion regulation strategy and trait anxiety to nodal parameters of the functional network, with the latter phenomenon additionally related to the global characteristics of the structural network.

As for the functional graph, the lack of associations with global parameters suggests that the behavioural differences might be related to alterations in the particular cognitive processes governed by specific brain areas rather than differential functioning of the entire network. This notion is supported by the fact that the regions identified here form part of the network involved in semantic and social cognition, with the latter process drawing on semantics-related computations (Zahn et al., 2007; Lambon Ralph et al., 2016; Jackson, 2021; Diveica et al., 2021). As such, it is not surprising that the very same areas have been implicated in cognitive emotion regulation (Brandl et al., 2019), mapping onto a neural network responsible for reinterpretation/reappraisal and distancing/perspective taking strategies (Morawetz et al., 2017). Taking into account the above evidence, our study suggests that alterations in the functional properties of the regions belonging to this particular circuitry may serve as the biological underpinning linking self-blaming behaviour and anxiety. The subsequent parts of the discussion will now go into details regarding the exact roles attributed to each area.

The left sATL, together with its contralateral homologue, is believed to be the brain’s locus for social semantic knowledge (Zahn et al., 2007; Rice et al., 2015; Binney et al., 2016; Olson et al., 2013). The region’s prominent role in self-blame has been pinpointed by studies showing the aberrant connectivity of this area with SCSR during guilt processing in MDD patients (Lythe et al., 2015; Green et al., 2012). By reporting sATL’s associations with both self-blaming behaviour and trait anxiety, our study underlines the key role it plays in the studied circuitry, implicating that the differential functional interactions of this region with the fronto-limbic emotional networks could underlie the prevalence of maladaptive forms of guilt observed across both mood and anxiety disorders (Kim et al., 2011; Candea and Szentagotai-Tătar, 2018; Green et al., 2012; González-García and Visser, 2023). The stronger interconnected left sATL in individuals with increased self-blame and trait-anxiety might reflect the tendency to ruminate over related negative social situations, which would require the sATL to access the related social-semantic information. The ATL has been described as a hub region that interconnects modality specific regions to obtain semantic (including social) information. The more frequently semantic items are processed, the stronger the connections become, meaning that less activity is required in the network to obtain the information (Lambon Ralph et al., 2016; Rogers et al., 2004; Rogers and McClelland, 2004). Therefore, a stronger interconnected ATL, could also possibly explain the decreased spontaneous brain activity in the right sATL in anxiety disorders (Wang et al., 2022). As such, our findings align with the idea that the sATL ‘communicates’ more effectively with other semantic-related regions in individuals with higher trait-anxiety and self-blame. The currently held view states that the sATL transmits the social information via the uncinate fasciculus to the medial prefrontal cortex, which in turn integrates it with emotional processes to regulate social behaviour (Sakata et al., 2019; Arioli et al., 2021). In the case of our study, the left sATL node was structurally connected to the ipsilateral SCSR in only 38.15% of the participants (Supplementary Figure 2). Nevertheless, we believed this finding to be related to the relatively small size of the node. Indeed, when larger ROIs were used, resembling the more widespread task fMRI activations, the fibres belonging to the tract were successfully identified in 85.52% of the subjects (Supplementary Figure 15), confirming that the sATL node indeed belongs to this functionally relevant circuitry.

The left IFGpo and adjacent parts of insula are the key elements of the brain system responsible for semantic control, i.e. the ability to access and manipulate meaningful information for example by inhibiting the dominant or amplifying the less dominant aspects of concepts and resolution of ambiguity or incongruent meanings (Jefferies, 2013; Jackson, 2021). Given the previously mentioned association between social and semantic cognition, it is not surprising that both regions are also implicated in processing moral information, with IFGpo involved in moral reasoning and anterior insular activity related to moral disgust and advantageous inequity (Diveica et al., 2021; Ying et al., 2018; Gao et al., 2018). Previous neuroimaging research has linked the structure of these areas with anxiety and guilt (Belden et al., 2014; Shang et al., 2014; Hu and Dolcos, 2017). Furthermore, in individuals with social anxiety disorder (SAD) the cognitive reappraisal network, which includes the left IFG as one of its nodes, exhibits decreased influence on the emotional reactivity network during reappraisal of interpersonal criticism (Jacob et al., 2019). Additionally, patients with obsessive-compulsive disorder (OCD) show decreased activity of the insula when experiencing deontological guilt, i.e. when one feels they have violated their own moral code (Basile et al., 2014). Our finding of a negative correlation between trait-anxiety and the efficiency of information integration around these two areas extends the previous literature by showing that the functional characteristics present in anxiety patients are also observed in subclinical groups, which is in line with the spectral nature of mood and anxiety disorders, and suggests that similar alterations in the architecture of brain networks may contribute to the guilt symptomatology across both subclinical and clinical disorder levels. As such, we interpret the functional disconnection of the left IFGpo and adjacent parts of insula reported here as a correlate of the deficiency in one’s ability to modulate the retrieval and selection of conceptual-level knowledge, diminishing the likelihood of successful regulation of social experiences labelled as guilt.

As for the structural graph, associations between anxiety and solely global parameters were found, indicating that, unlike in the functional network, anxiety is to a larger extent linked to the architecture of the circuitry as a whole rather than anatomical connections of its particular nodes. Given the high specificity of the areas included in the guilt network and the fact they belong to various canonical resting-state networks (Uddin et al., 2019), the results presented here are difficult to compare with the literature. Nevertheless, the connections between some of the areas included in the current circuitry were found to differ in the number of fibres between low and high trait anxiety individuals (Yang et al., 2020). The positive associations of the network’s clustering coefficient and anxiety scores at 10-15% densities indicate that in the more anxious participants we observe a neural architecture favouring increased local information processing. At the discussed sparsities, the guilt network appears to be organised into two submodules: the first one in the left temporal lobe, and the second consisting mainly of bilateral subcortical and cortical midline nodes (Supplementary Figures 16-17). As described before, these two subsystems are believed to be responsible, respectively, for the semantic and affective aspects of guilt experiences (Eslinger et al., 2021; Green et al., 2012; Lythe et al., 2015). Given that hypersensitivity to emotional stimuli has been reported both across anxiety patients and high-trait anxiety (Weinberg et al., 2010; Voegler et al., 2018; Hajcak et al., 2003), the increased communication within both modules could constitute a neural correlate of this characteristic and contribute to the prevalence of maladaptive forms of guilt observed in anxiety disorders (Candea and Szentagotai-Tătar, 2018).

One limitation of our study, however, is that the interpretation of the associations of anxiety and self-blaming behaviour with the characteristics of the guilt network remains limited only to the subclinical levels of anxiety. Before any potential clinical applications, the results reported here need to be confirmed in diagnosed individuals. Furthermore, on a more methodological note, we stress out the fact that the functional networks were constructed only using the significant positive correlations. While anticorrelations between brain regions are also known to be an important characteristic of a functional network’s architecture, this approach makes the topological graph measures more easily interpretable, and thus increases the utility of the reported findings (Qian et al, 2018). Last but not least, as the reconstruction of longer streamlines, including interhemispheric connections, is often less accurate, the contribution of the non-frontal right hemisphere regions to the structural connectome may be underestimated, while the contributions of the bilateral frontal regions may be biased in the opposite direction (Yeh et al., 2021). As such, the hubs of the structural network presented in the Supplementary Material should be treated with caution.

## 5. Conclusions

Recently, we have been observing a shift in paradigm towards personalised approaches in psychiatry with the hopes of increasing the efficiency of deployed treatments. Given the spectral nature of mood and anxiety disorders, our work maps onto this new framework by trying to find in a subclinical sample a link between a specific symptom (guilt) and the alterations in the underlying neural circuitry. Our findings provide an important insight that the prevalence of self-blame symptomatology in subclinical anxiety may be primarily related to the increased functional interconnectedness of the left sATL, which may manifest itself as a more pronounced communication of the guilt ‘label’ to its functionally coupled regions. This process may be further reinforced by the functional isolation of the left IFGpo and insula, a potential marker of deficiency in one’s ability to reinterpret the guilt-associated social experiences, and the network’s architecture favouring increased local information processing within the circuitrýs semantic and affective submodules.

## Funding

This publication forms part of several projects funded by the Spanish government: MCIN/AEI/10.13039/501100011033; grant RYC2019-028370-I and AEI PID2021-127516NB-I00; the Valencian Community: CIAICO/2021/088 and the University Jaume I: UJI-B2022-55.

## Data availability

The described analyses were performed with the use of an anonymised openly available dataset (Babayan et al., 2019).

## CRediT author statement

MRZ: Conceptualization, Formal analysis, Visualization, Writing - Original Draft. KB: Formal analysis, Writing - Review & Editing. VC: Conceptualization, Writing - Review & Editing. MV: Conceptualization, Supervision, Writing - Review & Editing.

## Disclosure statement

The authors report there are no competing interests to declare.

## Ethical statement

This work was performed using an anonymised openly available dataset (Babayan et al., 2019) and thus was not subjected to ethical committee approval.

## Notes

### Competing Interest Statement

The authors have declared no competing interest.

