## Supplementary Materials for "Graph analysis of the guilt network highlights associations with subclinical anxiety and self-blame"

**Supplementary Table 1.** Graph theory measures extracted from the functional and structural guilt networks.

| **Measure** | **Definition** |
|---|---|
| Strength | Sum of the edge weights assigned to the node (Barrat et al., 2004) |
| Weighted clustering coefficient | An extension of unweighted clustering coefficient, i.e. the proportion of the number of links between the node and its immediate neighbours divided by the number of links that could possibly exist between them, that takes into account the edge weights (Barrat et al., 2004). |
| Betweenness centrality | Number of the shortest paths in the graph that pass through the node (Freeman, 1979). |
| Local efficiency | Average of the inverse of the distances between the node's neighbours through the rest of the network (Latora and Marchiori, 2001). |
| Global efficiency | Average of the inverse of the distances between all pairs of nodes (Latora and Marchiori, 2001) |
| Modularity | Extent to which a graph can be divided into clearly separated communities (Traag et al., 2019). |

**Supplementary Table 2.** The hub characteristics of the nodes of the functional network across all the network densities (5-30%). The betweenness centrality was first normalised within each individual's network (1/N, where N is the highest observed betweenness centrality at the given sparsity level) and then averaged across all the participants. Two frontal regions, i.e. the left superior frontal gyrus and medial frontal gyrus (BA 10), and one temporal region (left middle temporal gyrus) appear to serve hub functions across all thresholding levels. In the parietal lobe the area displaying hub characteristics depends on the network density: with increasing sparsity it moves from the left precuneus to the left posterior cingulate cortex/parieto-occipital sulcus. The node-to-node comparisons were calculated with permutation t-tests (perm.t.test function from the MKinfer package in R) and were corrected for multiple comparisons using the Bonferroni method (FWE < 0.05). Their significance for each network density is presented in Supplementary Figures 3-8.

| **Location** | **Betweenness centrality** | | | | | |
|---|---|---|---|---|---|---|
| | **5%** | **10%** | **15%** | **20%** | **25%** | **30%** |
| R medial orbitofrontal cortex | 0.049 | 0.092 | 0.183 | 0.167 | 0.207 | 0.221 |
| R subgenual cingulate cortex and septal area | 0.032 | 0.094 | 0.112 | 0.142 | 0.160 | 0.193 |
| L subgenual cingulate cortex and septal area | 0.066 | 0.168 | 0.259 | 0.221 | 0.247 | 0.229 |
| L anterior cingulate cortex | 0.034 | 0.091 | 0.178 | 0.213 | 0.300 | 0.327 |
| L superior frontal gyrus | 0.317 | 0.319 | 0.354 | 0.401 | 0.418 | 0.444 |
| L medial frontal gyrus (BA 6) | 0.077 | 0.138 | 0.215 | 0.231 | 0.294 | 0.310 |
| L medial frontal gyrus (BA 10) | 0.260 | 0.389 | 0.447 | 0.429 | 0.443 | 0.464 |
| L medial frontal gyrus (BA 9) | 0.116 | 0.088 | 0.105 | 0.113 | 0.125 | 0.126 |
| L inferior frontal gyrus pars opercularis and insula | 0.065 | 0.077 | 0.131 | 0.158 | 0.170 | 0.195 |
| R inferior frontal gyrus pars opercularis | 0.038 | 0.080 | 0.173 | 0.241 | 0.215 | 0.198 |
| L superior temporal gyrus | 0.191 | 0.250 | 0.153 | 0.152 | 0.196 | 0.211 |
| L middle temporal gyrus | 0.309 | 0.380 | 0.372 | 0.430 | 0.443 | 0.454 |
| L superior anterior temporal lobe | 0.000 | 0.028 | 0.074 | 0.071 | 0.064 | 0.071 |
| R superior anterior temporal lobe | 0.013 | 0.079 | 0.147 | 0.185 | 0.250 | 0.259 |
| L posterior cingulate cortex and parieto-occipital sulcus | 0.143 | 0.271 | 0.427 | 0.462 | 0.504 | 0.517 |
| L precuneus | 0.270 | 0.311 | 0.248 | 0.226 | 0.264 | 0.285 |
| L supramarginal gyrus | 0.068 | 0.187 | 0.270 | 0.303 | 0.328 | 0.351 |
| L cuneus | 0.023 | 0.104 | 0.118 | 0.197 | 0.181 | 0.195 |
| R lingual gyrus | 0.056 | 0.100 | 0.167 | 0.220 | 0.226 | 0.230 |

Abbreviations: L, left; R, right; BA, Brodmann area.

**Supplementary Table 3.** The hub characteristics of the nodes of the structural network across all the network densities (5-30%). The betweenness centrality was first normalised within each individual's network (1/N, where N is the highest observed betweenness centrality at the given sparsity level) and then averaged across all the participants. The left anterior cingulate cortex and medial frontal gyrus (BA 10) functioned as hubs across all network densities. At higher sparsities, additional hubs emerged in the left medial frontal gyrus (BA 9), inferior frontal gyrus pars opercularis and insula, and supramarginal gyrus. The node-to-node comparisons were calculated with permutation t-tests (perm.t.test function from the MKinfer package in R) and were corrected for multiple comparisons using the Bonferroni method (FWE < 0.05). Their significance for each network density is presented in Supplementary Figures 9-14.

| **Location** | **Betweenness centrality** | | | | | |
|---|---|---|---|---|---|---|
| | **5%** | **10%** | **15%** | **20%** | **25%** | **30%** |
| R medial orbitofrontal cortex | 0.044 | 0.123 | 0.065 | 0.062 | 0.059 | 0.059 |
| R subgenual cingulate cortex and septal area | 0.291 | 0.165 | 0.078 | 0.076 | 0.072 | 0.078 |
| L subgenual cingulate cortex and septal area | 0.169 | 0.329 | 0.213 | 0.203 | 0.192 | 0.195 |
| L anterior cingulate cortex | 0.329 | 0.788 | 0.893 | 0.919 | 0.927 | 0.934 |
| L superior frontal gyrus | 0.606 | 0.275 | 0.257 | 0.280 | 0.270 | 0.265 |
| L medial frontal gyrus (BA 6) | 0.000 | 0.121 | 0.292 | 0.333 | 0.335 | 0.335 |
| L medial frontal gyrus (BA 10) | 0.817 | 0.738 | 0.547 | 0.534 | 0.515 | 0.511 |
| L medial frontal gyrus (BA 9) | 0.090 | 0.138 | 0.437 | 0.527 | 0.533 | 0.524 |
| L inferior frontal gyrus pars opercularis and insula | 0.000 | 0.125 | 0.535 | 0.663 | 0.646 | 0.638 |
| R inferior frontal gyrus pars opercularis | 0.000 | 0.000 | 0.000 | 0.000 | 0.005 | 0.005 |
| L superior temporal gyrus | 0.000 | 0.000 | 0.004 | 0.010 | 0.010 | 0.009 |
| L middle temporal gyrus | 0.049 | 0.130 | 0.250 | 0.287 | 0.278 | 0.274 |
| L superior anterior temporal lobe | 0.000 | 0.017 | 0.016 | 0.024 | 0.022 | 0.022 |
| R superior anterior temporal lobe | 0.000 | 0.000 | 0.000 | 0.000 | 0.000 | 0.003 |
| L posterior cingulate cortex and parieto-occipital sulcus | 0.000 | 0.056 | 0.226 | 0.275 | 0.299 | 0.337 |
| L precuneus | 0.009 | 0.114 | 0.280 | 0.317 | 0.331 | 0.345 |
| L supramarginal gyrus | 0.046 | 0.163 | 0.438 | 0.536 | 0.520 | 0.513 |
| L cuneus | 0.000 | 0.014 | 0.033 | 0.058 | 0.080 | 0.102 |
| R lingual gyrus | 0.000 | 0.000 | 0.000 | 0.000 | 0.009 | 0.026 |

Abbreviations: L, left; R, right; BA, Brodmann area.

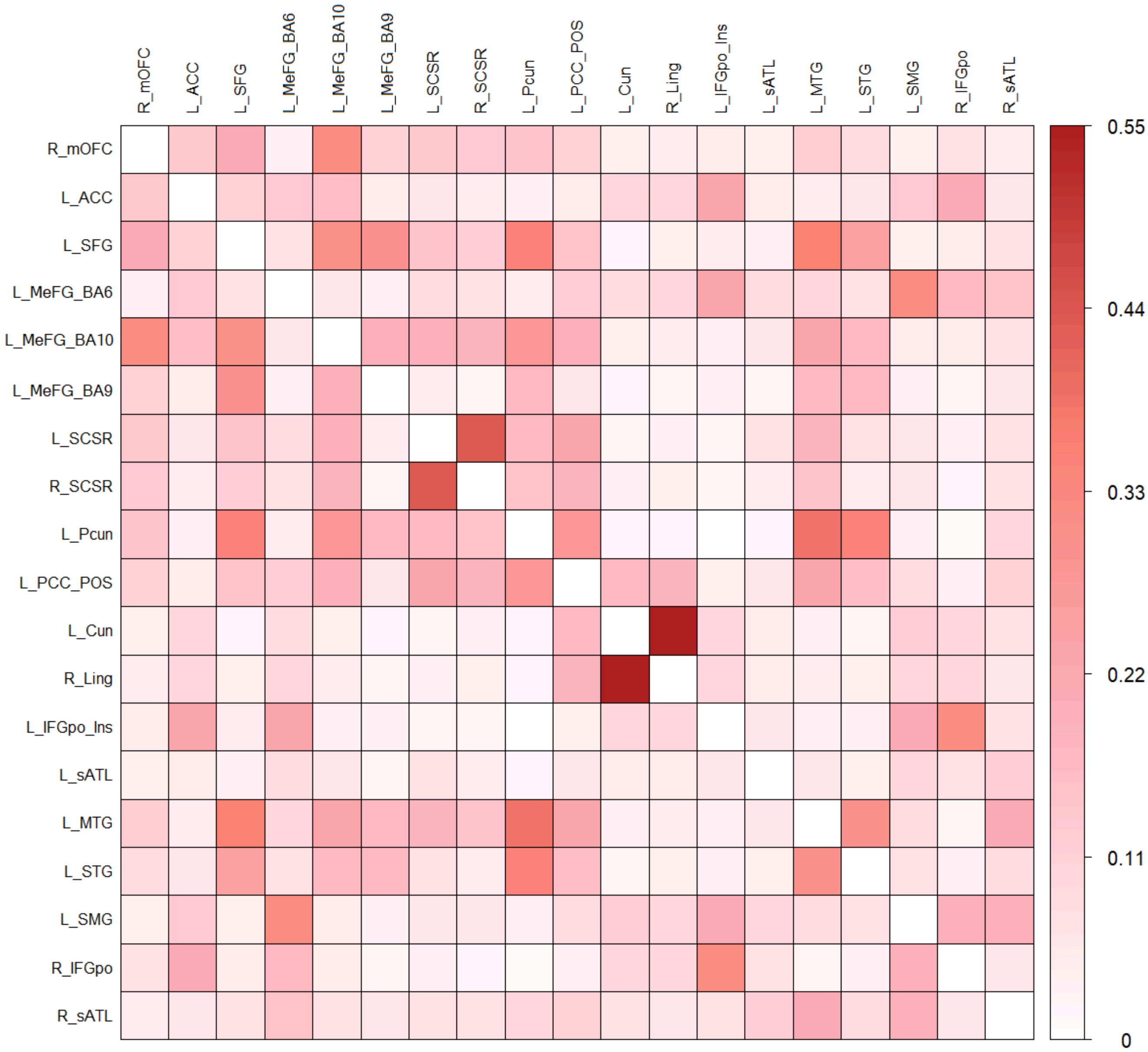


**Supplementary Figure 1.** Group-averaged functional connectivity between the nodes of the guilt network. Abbreviations: R_mOFC, right medial orbitofrontal cortex; L_ACC, left anterior cingulate cortex; L_SFG, left superior frontal gyrus; L_MeFG_BA6, left medial frontal gyrus (Brodmann area 6); L_MeFG_BA10, left medial frontal gyrus (Brodmann area 10); L_MeFG_BA9, left medial frontal gyrus (Brodmann area 9); L_SCSR, left subgenual cingulate cortex and adjacent septal area; R_SCSR, right subgenual cingulate cortex and adjacent septal area; L_Pcun, left precuneus; L_PCC_POS, left posterior cingulate cortex and parieto-occipital sulcus; L_Cun, left cuneus; R_Ling, right lingual gyrus; L_IFGpo_Ins, left inferior frontal gyrus pars opercularis and insula; L_sATL, left superior anterior temporal lobe; L_MTG, left middle temporal gyrus; L_STG, left superior temporal gyrus; L_SMG, left supramarginal gyrus; R_IFGpo, right inferior frontal gyrus pars opercularis; R_sATL, right superior anterior temporal lobe.

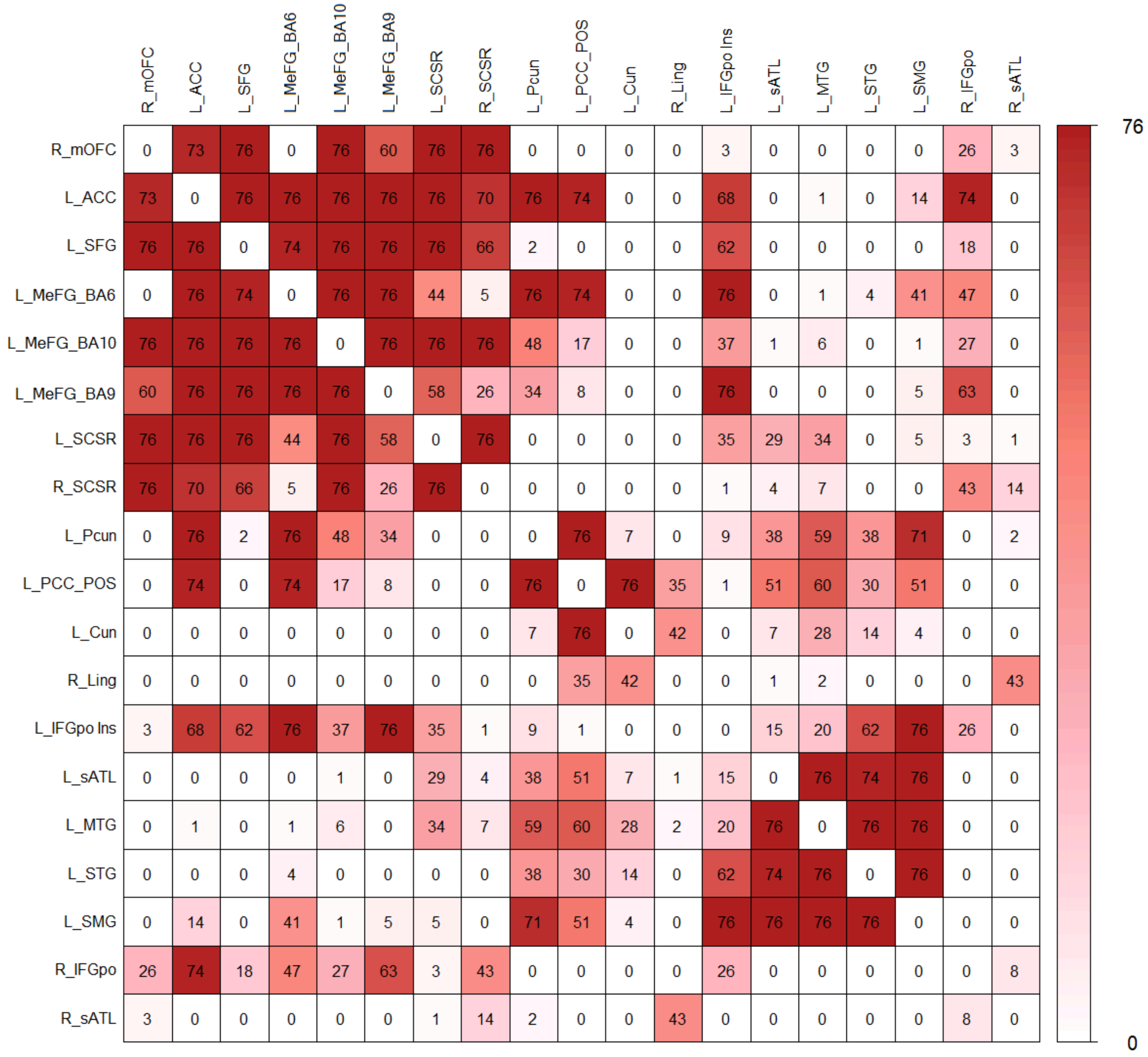


**Supplementary Figure 2.** Structural connectivity between the nodes of the unthresholded guilt network. The numbers reflect how many individuals were found to have a given connection. Abbreviations: R_mOFC, right medial orbitofrontal cortex; L_ACC, left anterior cingulate cortex; L_SFG, left superior frontal gyrus; L_MeFG_BA6, left medial frontal gyrus (Brodmann area 6); L_MeFG_BA10, left medial frontal gyrus (Brodmann area 10); L_MeFG_BA9, left medial frontal gyrus (Brodmann area 9); L_SCSR, left subgenual cingulate cortex and adjacent septal area; R_SCSR, right subgenual cingulate cortex and adjacent septal area; L_Pcun, left precuneus; L_PCC_POS, left posterior cingulate cortex and parieto-occipital sulcus; L_Cun, left cuneus; R_Ling, right lingual gyrus; L_IFGpo_Ins, left inferior frontal gyrus pars opercularis and insula; L_sATL, left superior anterior temporal lobe; L_MTG, left middle temporal gyrus; L_STG, left superior temporal gyrus; L_SMG, left supramarginal gyrus; R_IFGpo, right inferior frontal gyrus pars opercularis; R_sATL, right superior anterior temporal lobe.

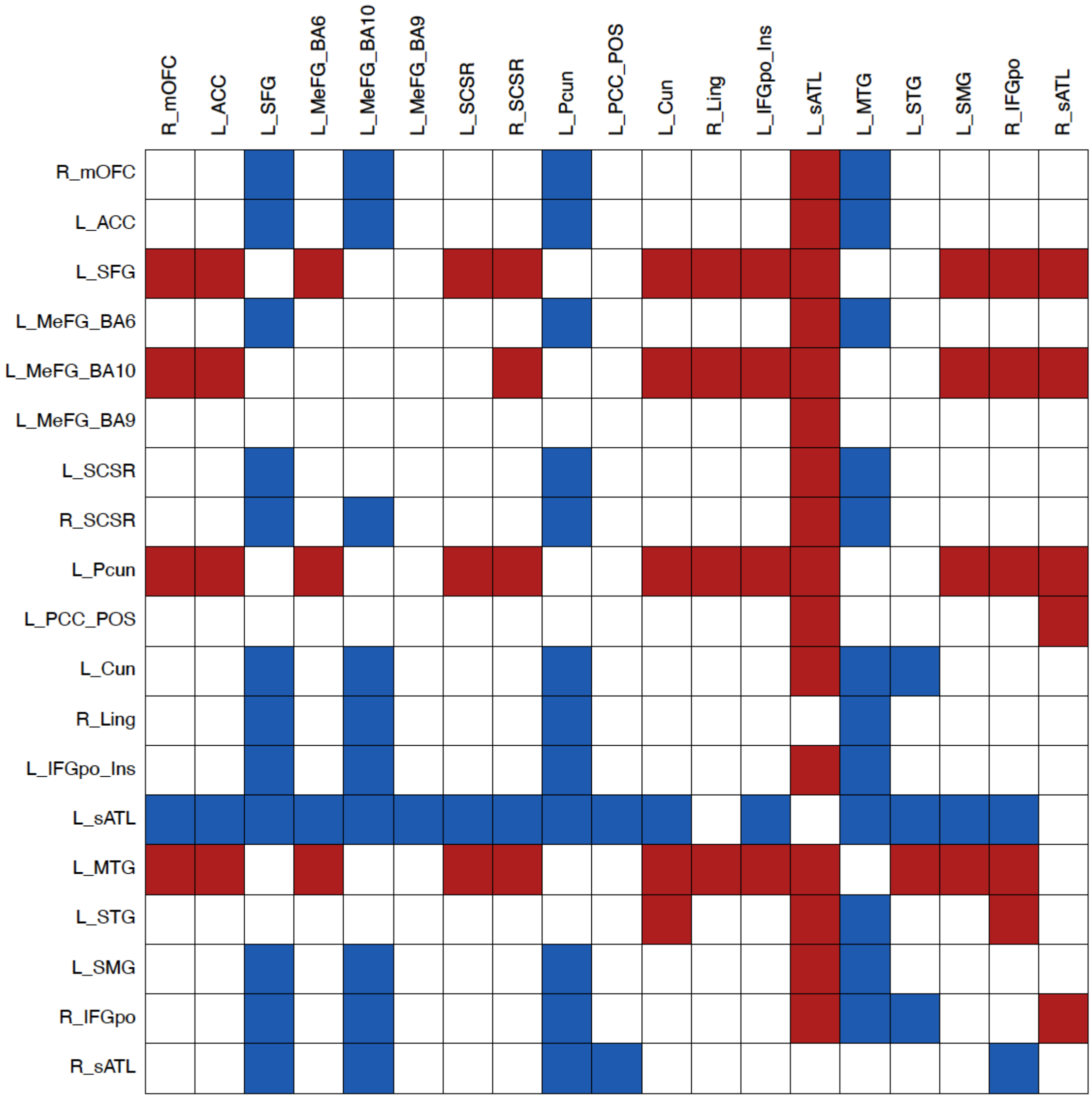


**Supplementary Figure 3.** The nodal comparisons in the hub characteristics for the functional network at 5% density. The red colour indicates significantly higher (FWE < 0.05) normalised betweenness centrality for the region in the y axis of the matrix, while the blue colour indicates the opposite.

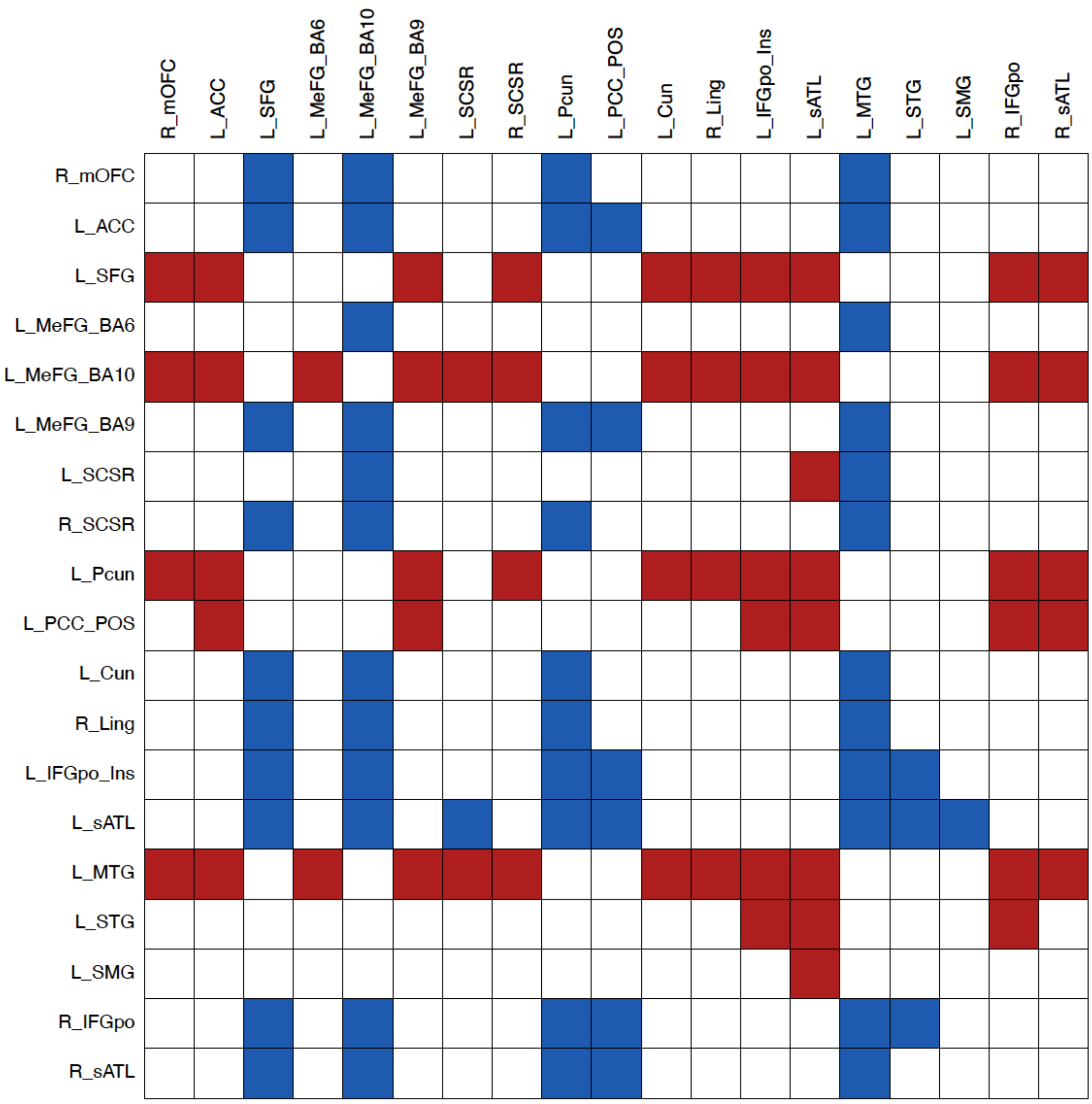


**Supplementary Figure 4.** The nodal comparisons in the hub characteristics for the functional network at 10% density. The red colour indicates significantly higher (FWE < 0.05) normalised betweenness centrality for the region in the y axis of the matrix, while the blue colour indicates the opposite.

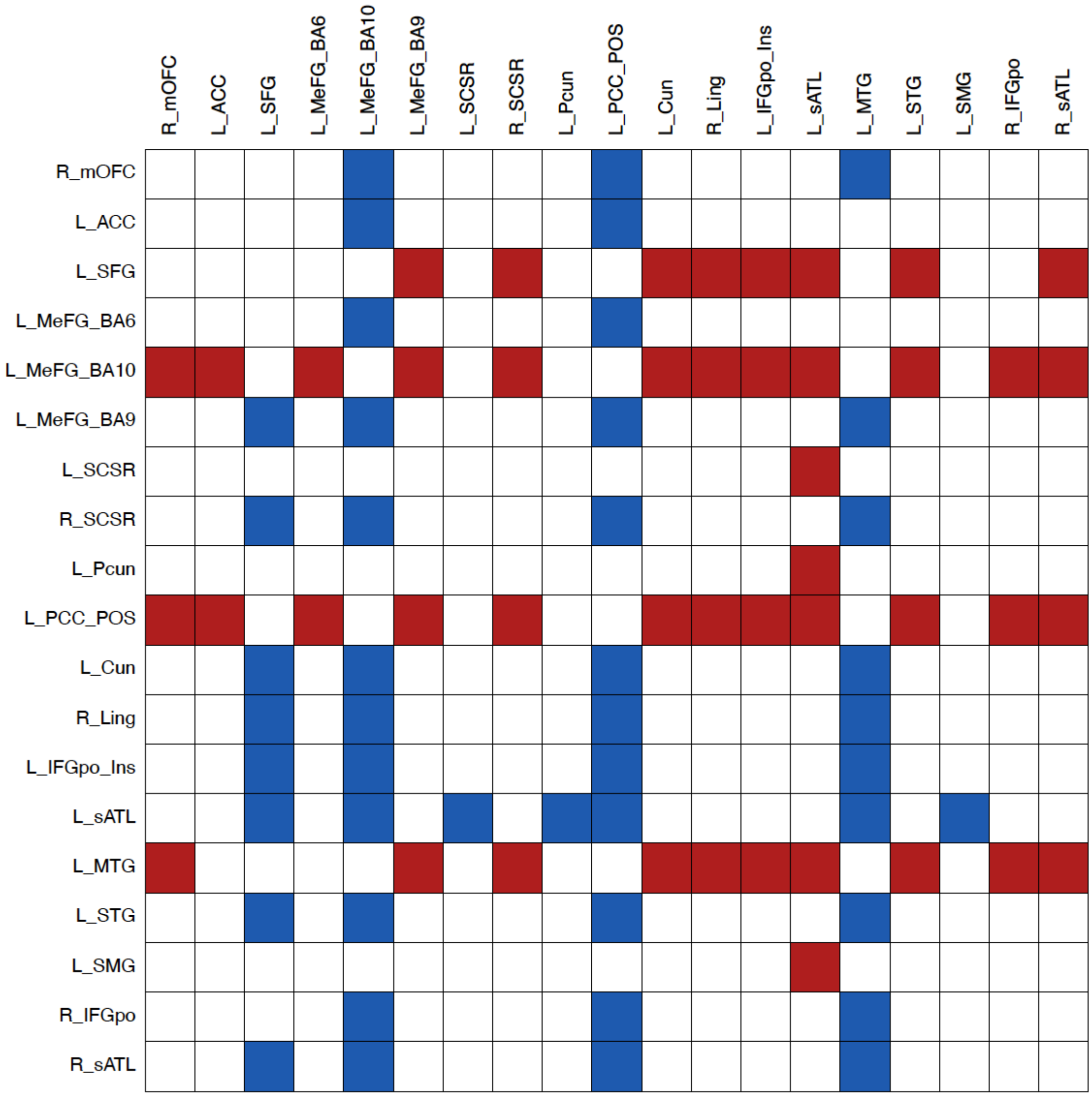


**Supplementary Figure 5.** The nodal comparisons in the hub characteristics for the functional network at 15% density. The red colour indicates significantly higher (FWE < 0.05) normalised betweenness centrality for the region in the y axis of the matrix, while the blue colour indicates the opposite.

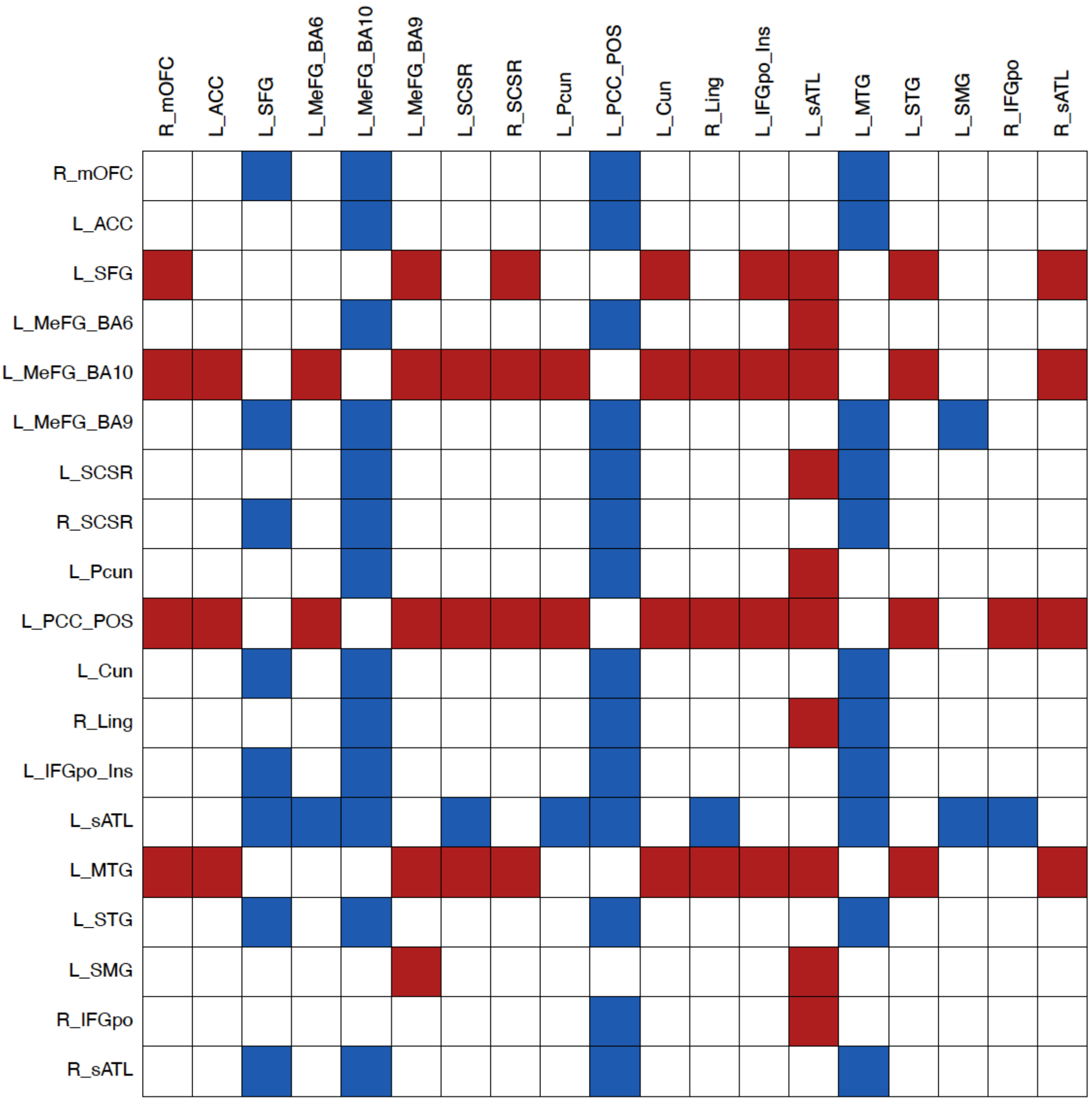


**Supplementary Figure 6.** The nodal comparisons in the hub characteristics for the functional network at 20% density. The red colour indicates significantly higher (FWE < 0.05) normalised betweenness centrality for the region in the y axis of the matrix, while the blue colour indicates the opposite.

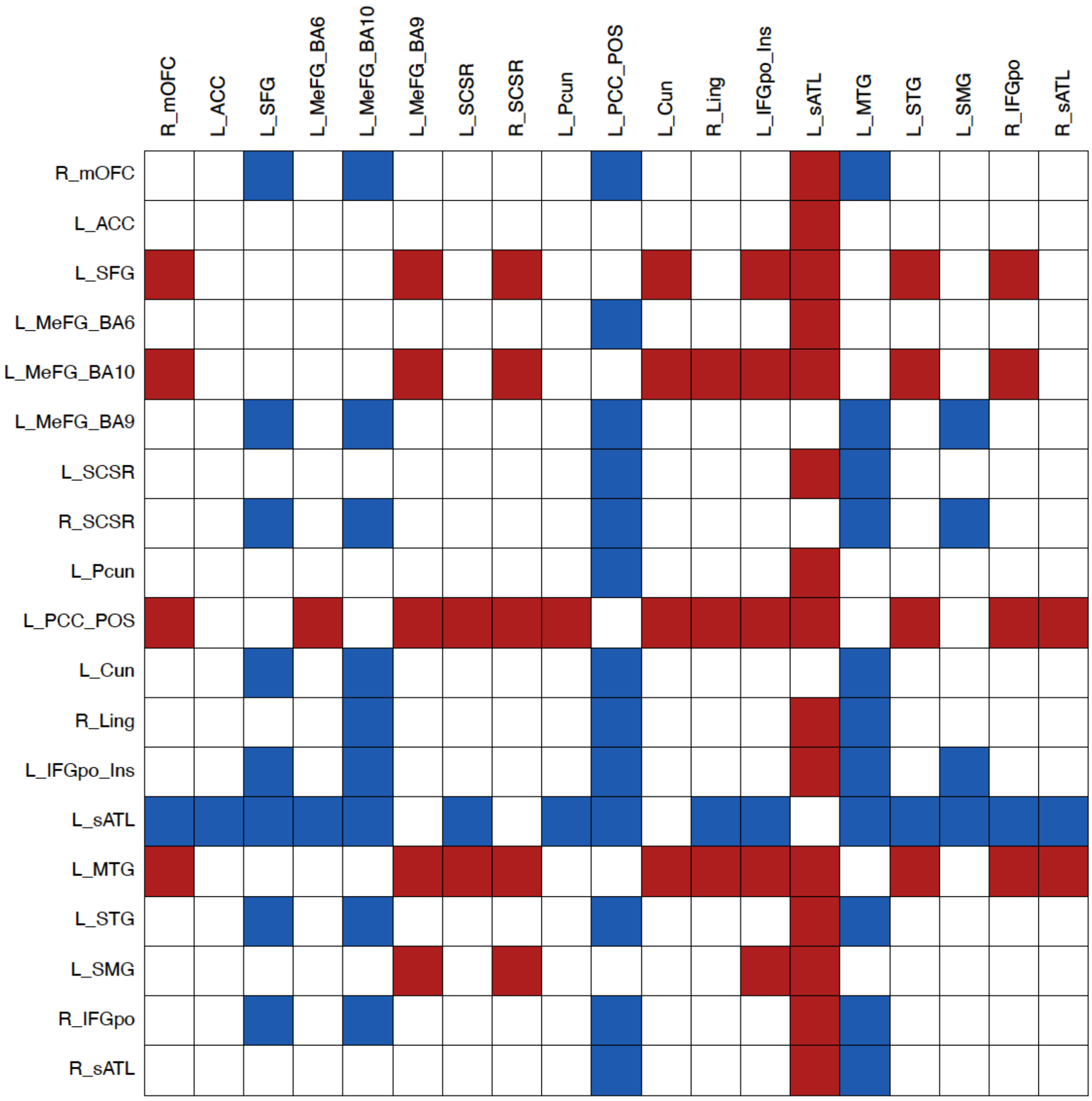


**Supplementary Figure 7.** The nodal comparisons in the hub characteristics for the functional network at 25% density. The red colour indicates significantly higher (FWE < 0.05) normalised betweenness centrality for the region in the y axis of the matrix, while the blue colour indicates the opposite.

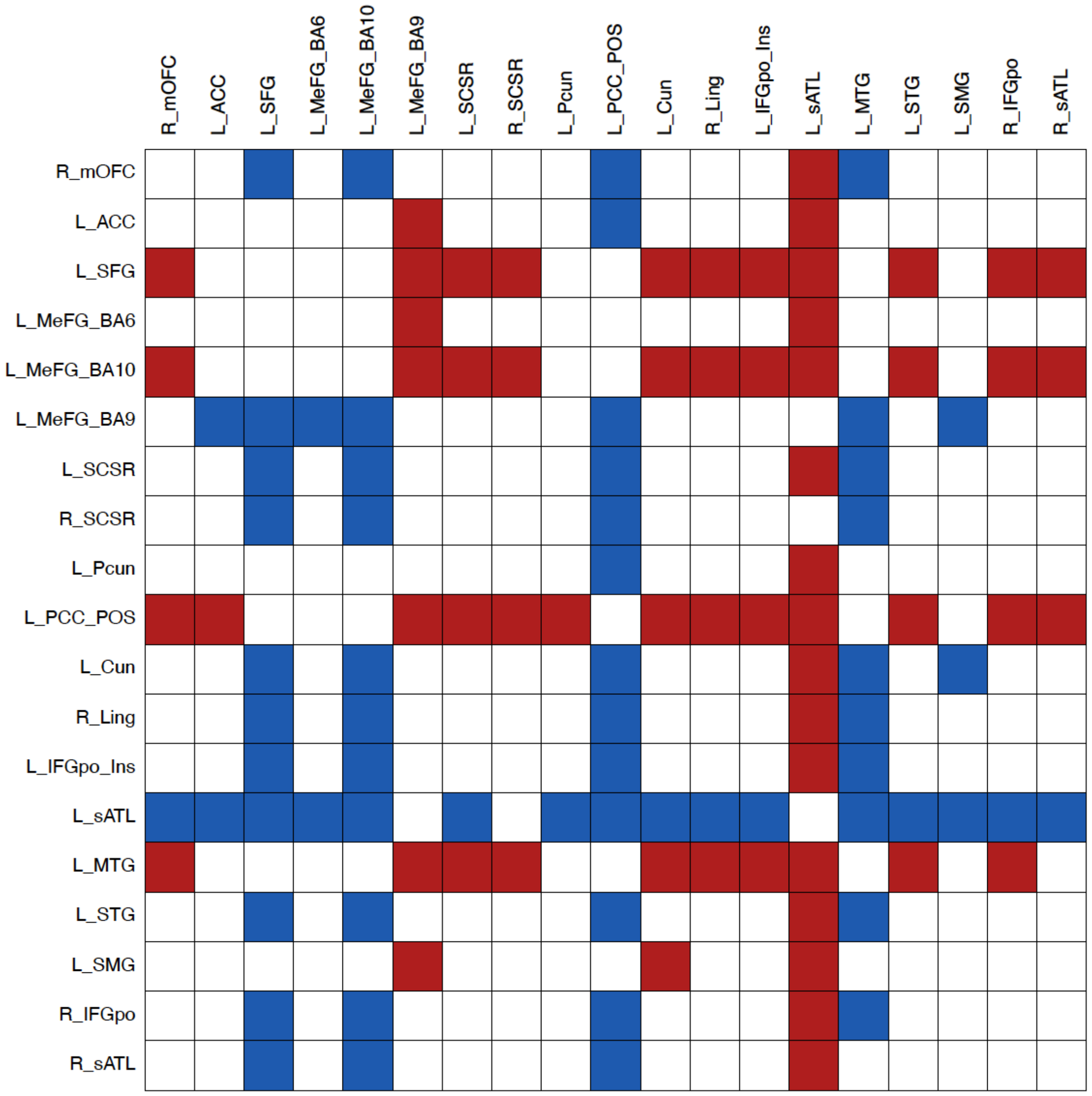


**Supplementary Figure 8.** The nodal comparisons in the hub characteristics for the functional network at 30% density. The red colour indicates significantly higher (FWE < 0.05) normalised betweenness centrality for the region in the y axis of the matrix, while the blue colour indicates the opposite.

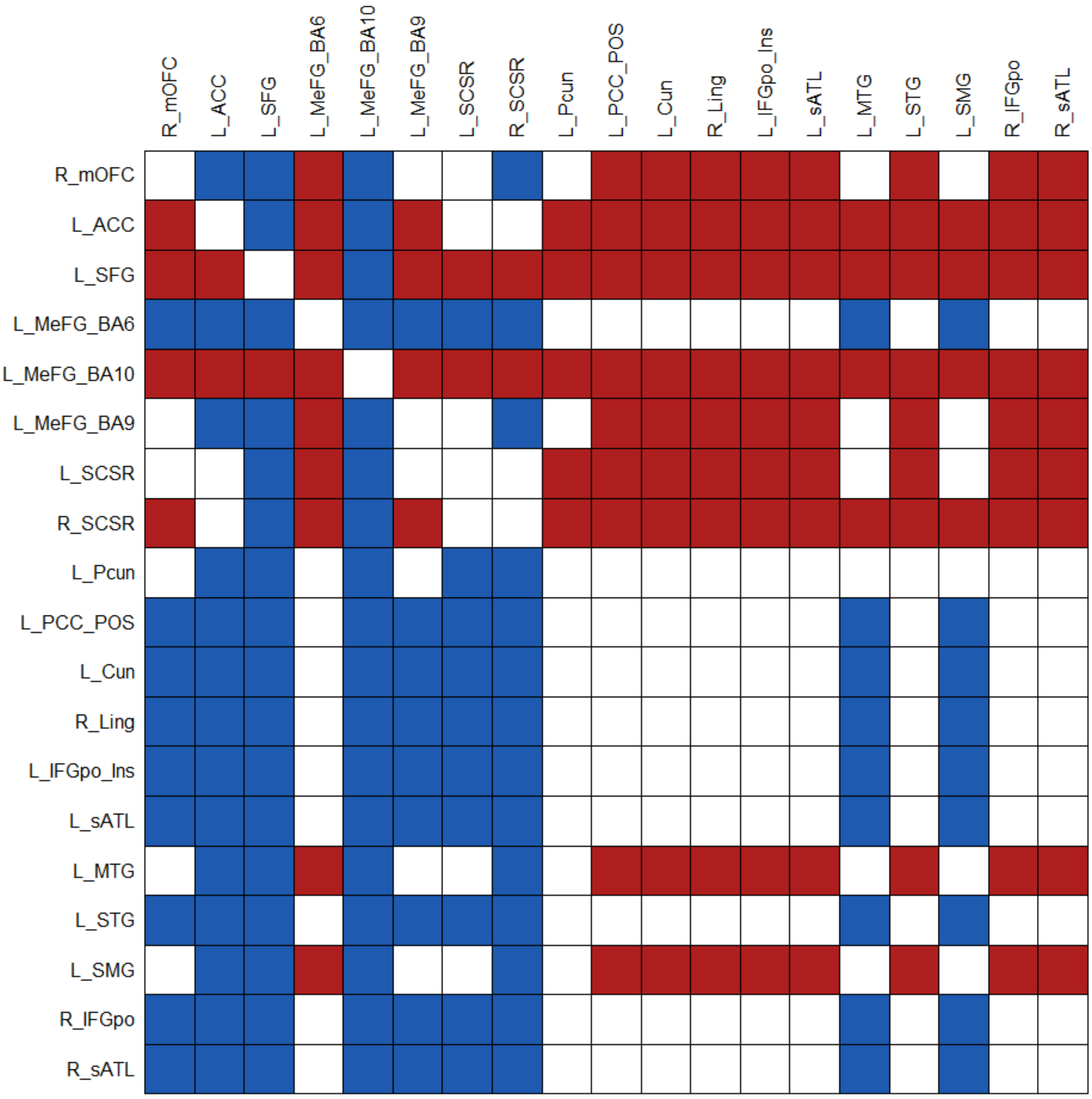


**Supplementary Figure 9.** The nodal comparisons in the hub characteristics for the structural network at 5% density. The red colour indicates significantly higher (FWE < 0.05) normalised betweenness centrality for the region in the y axis of the matrix, while the blue colour indicates the opposite.

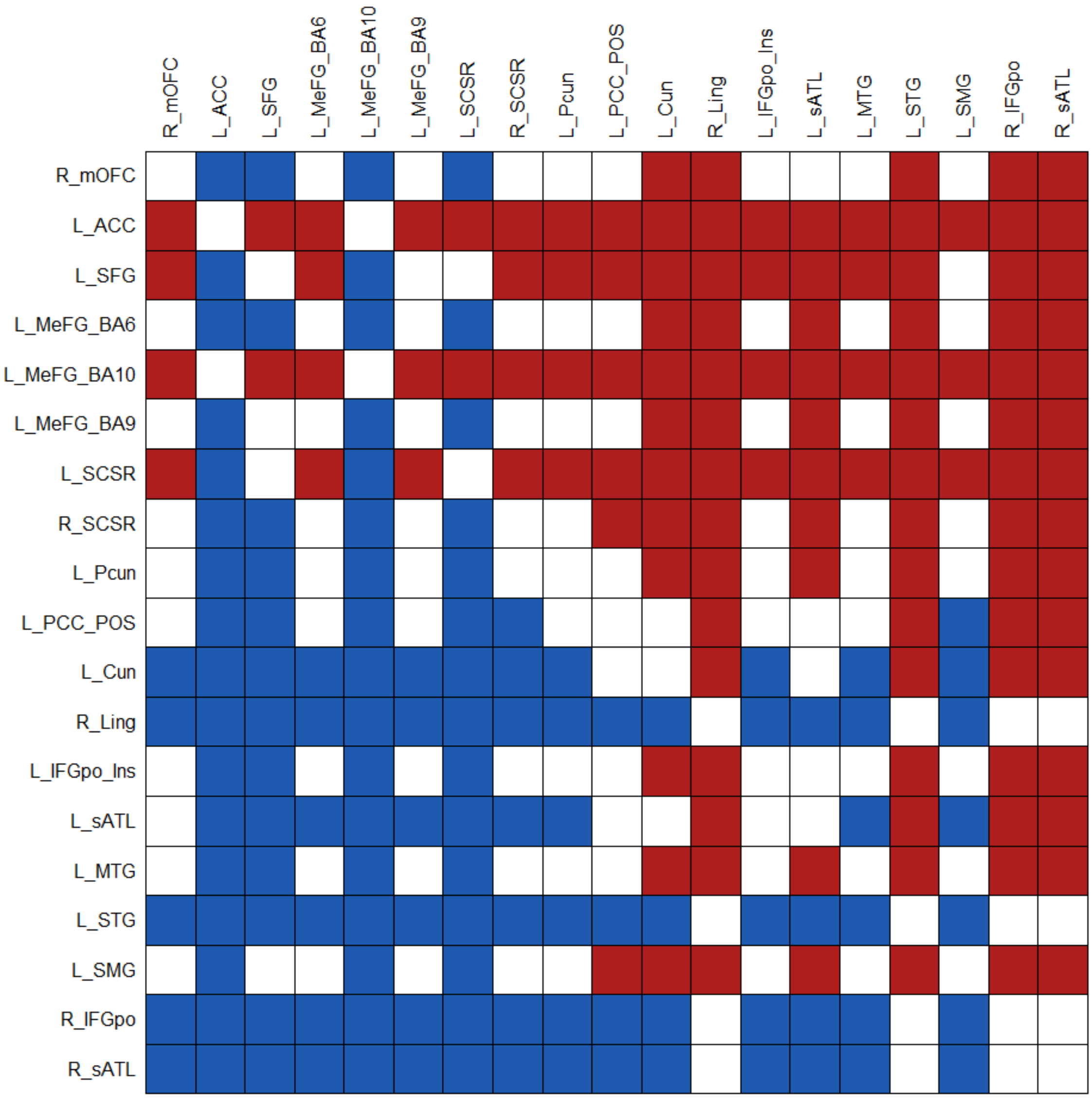


**Supplementary Figure 10.** The nodal comparisons in the hub characteristics for the structural network at 10% density. The red colour indicates significantly higher (FWE < 0.05) normalised betweenness centrality for the region in the y axis of the matrix, while the blue colour indicates the opposite.

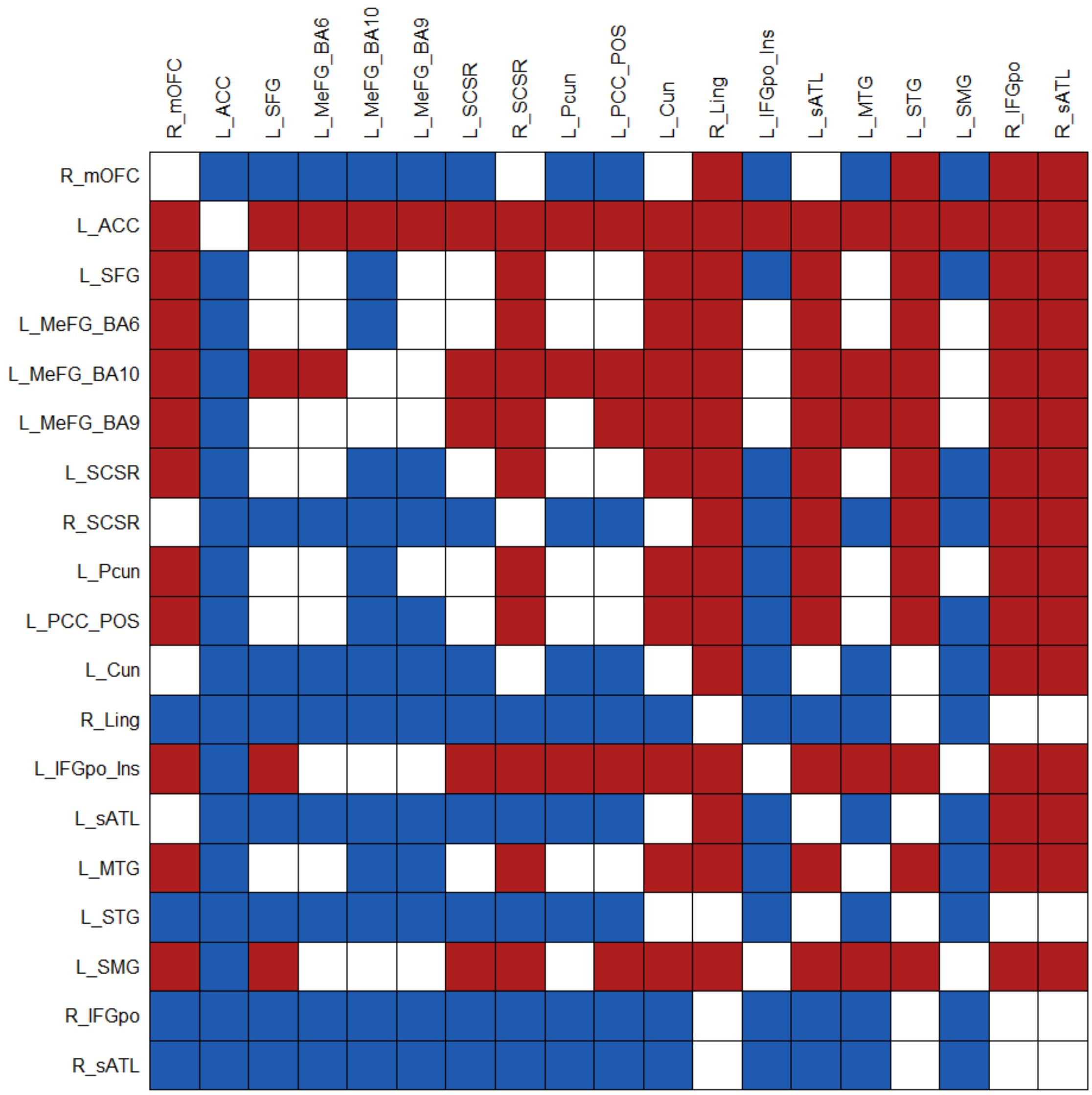


**Supplementary Figure 11.** The nodal comparisons in the hub characteristics for the structural network at 15% density. The red colour indicates significantly higher (FWE < 0.05) normalised betweenness centrality for the region in the y axis of the matrix, while the blue colour indicates the opposite.

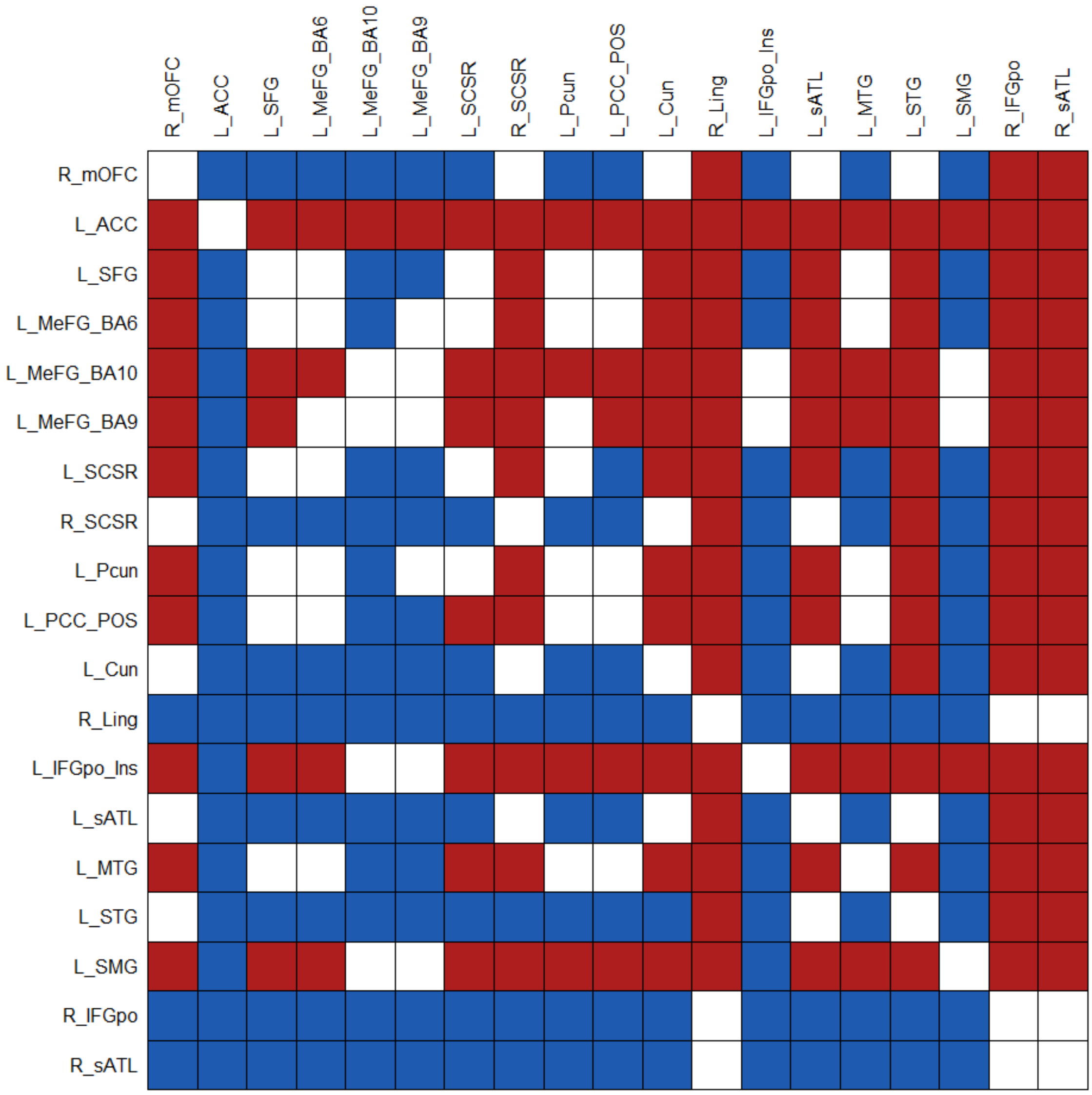


**Supplementary Figure 12.** The nodal comparisons in the hub characteristics for the structural network at 20% density. The red colour indicates significantly higher (FWE < 0.05) normalised betweenness centrality for the region in the y axis of the matrix, while the blue colour indicates the opposite.

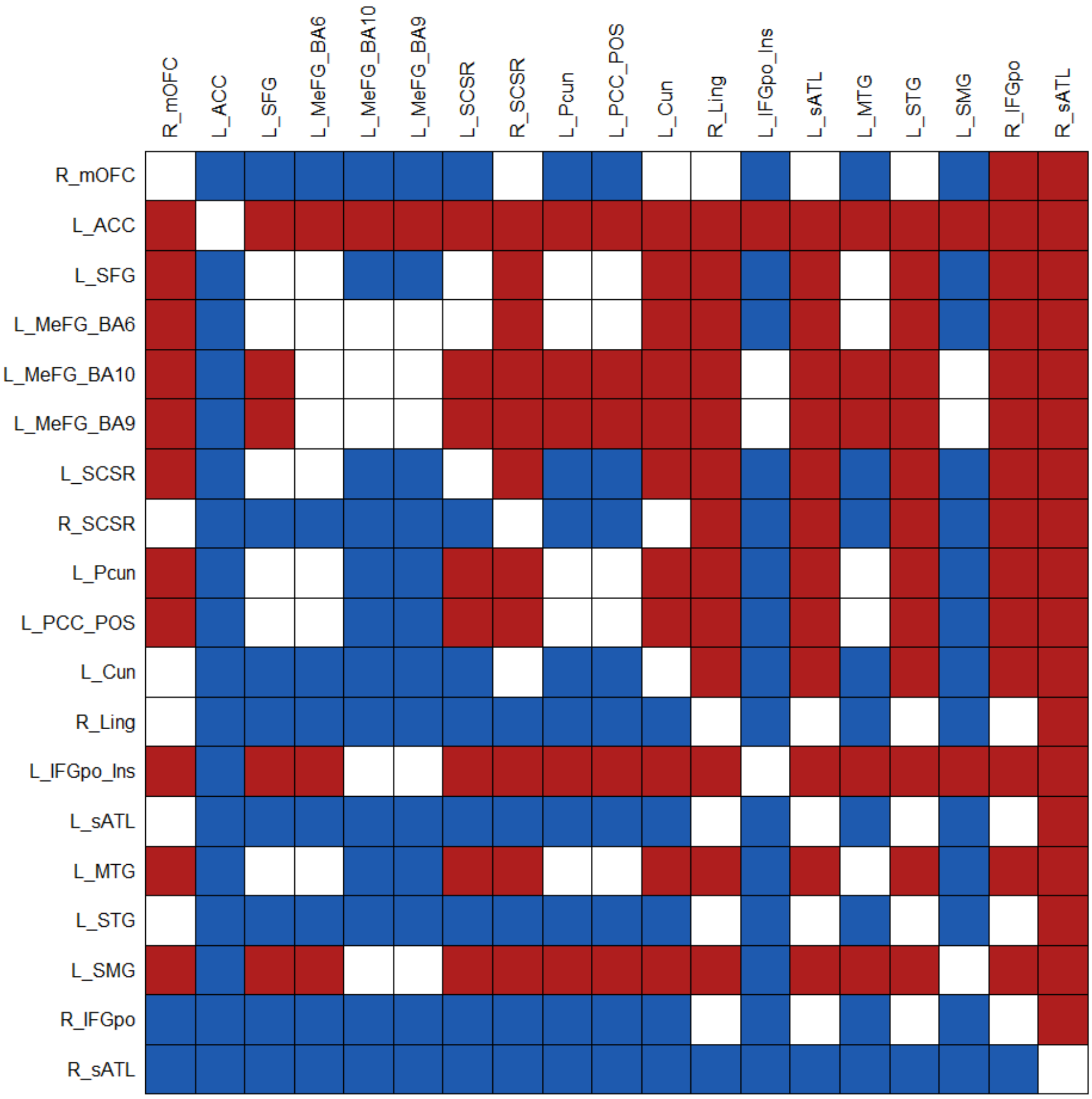


**Supplementary Figure 13.** The nodal comparisons in the hub characteristics for the structural network at 25% density. The red colour indicates significantly higher (FWE < 0.05) normalised betweenness centrality for the region in the y axis of the matrix, while the blue colour indicates the opposite.

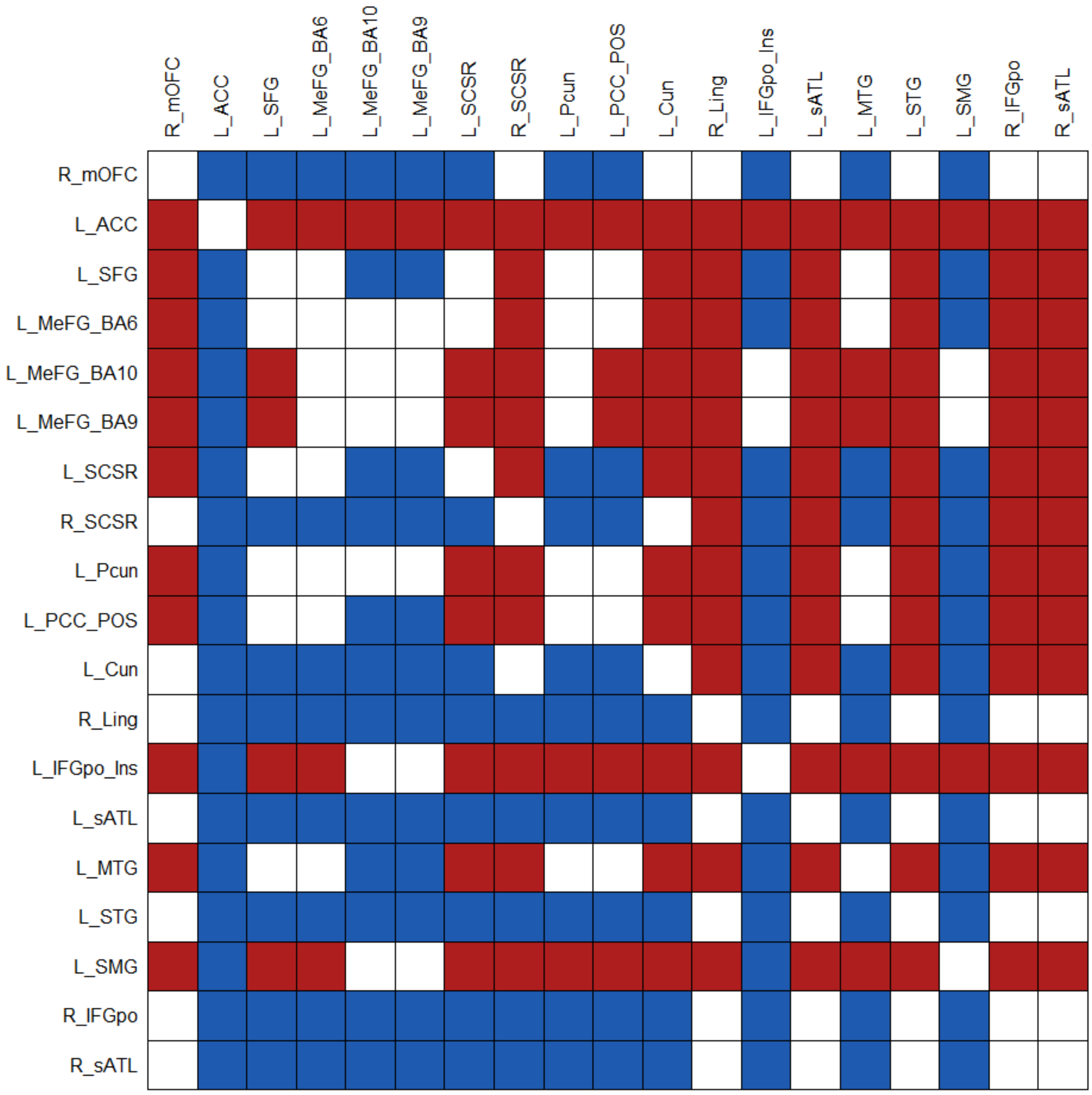


**Supplementary Figure 14.** The nodal comparisons in the hub characteristics for the structural network at 30% density. The red colour indicates significantly higher (FWE < 0.05) normalised betweenness centrality for the region in the y axis of the matrix, while the blue colour indicates the opposite.

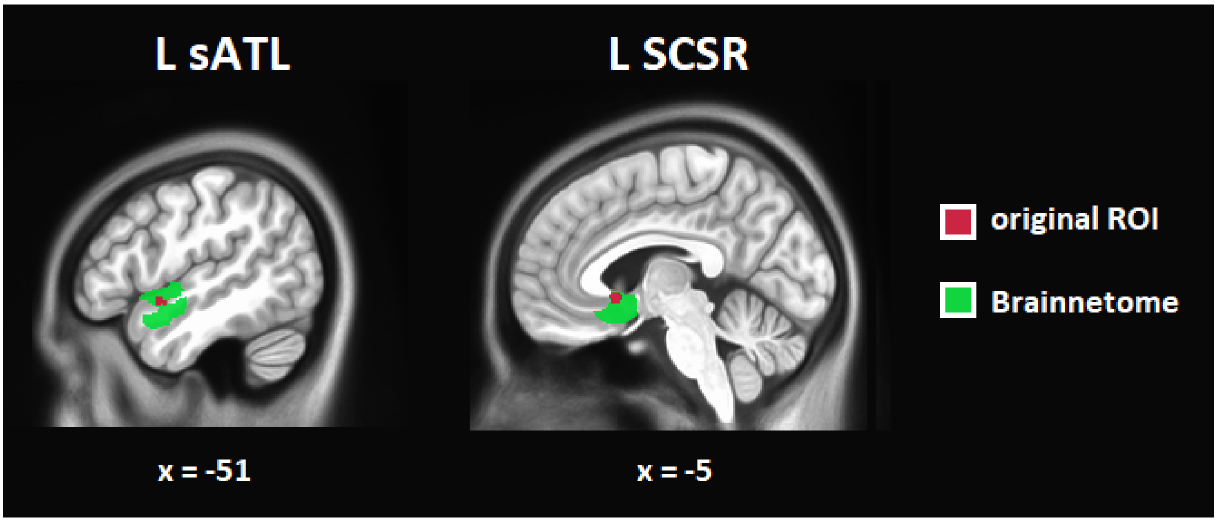


**Supplementary Figure 15.** Regions of interests (ROIs) used for investigating structural connectivity between the left superior anterior temporal lobe (L sATL) and the left subgenual cingulate cortex and adjacent septal area (L SCSR). Probabilistic tractography between the original ROIs (purple) identified uncinate fasciculus fibres linking these two structures only in 38.15% of the participants. To assess whether this finding was related to the particularly small size of the nodes, the analysis was repeated using the areas from the Brainnetome atlas that the original ROIs mapped onto (green). Indeed, in the latter case the fibres connecting the two regions were identified in 85.52% of the participants, confirming the presence of anatomical connectivity between L sATL and L SCSR.

| | R_mOFC | L_ACC | L_SFG | L_MeFG_BA6 | L_MeFG_BA10 | L_MeFG_BA9 | L_SCSR | R_SCSR | L_Pcun | L_PCC_POS | L_Cun | R_Ling | L_IFGpo_Ins | L_sATL | L_MTG | L_STG | L_SMG | R_IFGpo | R_sATL |
|---|---|---|---|---|---|---|---|---|---|---|---|---|---|---|---|---|---|---|---|
| R_mOFC | 0 | 0 | 42 | 0 | 65 | 0 | 34 | 73 | 0 | 0 | 0 | 0 | 0 | 0 | 0 | 0 | 0 | 0 | 0 |
| L_ACC | 0 | 0 | 76 | 51 | 76 | 76 | 61 | 0 | 37 | 6 | 0 | 0 | 3 | 0 | 0 | 0 | 0 | 2 | 0 |
| L_SFG | 42 | 76 | 0 | 0 | 76 | 76 | 0 | 0 | 0 | 0 | 0 | 0 | 0 | 0 | 0 | 0 | 0 | 0 | 0 |
| L_MeFG_BA6 | 0 | 51 | 0 | 0 | 9 | 5 | 0 | 0 | 43 | 0 | 0 | 0 | 6 | 0 | 0 | 0 | 0 | 0 | 0 |
| L_MeFG_BA10 | 65 | 76 | 76 | 9 | 0 | 75 | 36 | 2 | 1 | 0 | 0 | 0 | 0 | 0 | 0 | 0 | 0 | 0 | 0 |
| L_MeFG_BA9 | 0 | 76 | 76 | 5 | 75 | 0 | 0 | 0 | 0 | 0 | 0 | 0 | 14 | 0 | 0 | 0 | 0 | 2 | 0 |
| L_SCSR | 34 | 61 | 0 | 0 | 36 | 0 | 0 | 76 | 0 | 0 | 0 | 0 | 0 | 0 | 0 | 0 | 0 | 0 | 0 |
| R_SCSR | 73 | 0 | 0 | 0 | 2 | 0 | 76 | 0 | 0 | 0 | 0 | 0 | 0 | 0 | 0 | 0 | 0 | 0 | 0 |
| L_Pcun | 0 | 37 | 0 | 43 | 1 | 0 | 0 | 0 | 0 | 30 | 0 | 0 | 0 | 0 | 0 | 0 | 0 | 0 | 0 |
| L_PCC_POS | 0 | 6 | 0 | 0 | 0 | 0 | 0 | 0 | 30 | 0 | 60 | 0 | 0 | 0 | 0 | 0 | 0 | 0 | 0 |
| L_Cun | 0 | 0 | 0 | 0 | 0 | 0 | 0 | 0 | 0 | 60 | 0 | 12 | 0 | 0 | 0 | 0 | 0 | 0 | 0 |
| R_Ling | 0 | 0 | 0 | 0 | 0 | 0 | 0 | 0 | 0 | 0 | 12 | 0 | 0 | 0 | 0 | 0 | 0 | 0 | 0 |
| L_IFGpo_Ins | 0 | 3 | 0 | 6 | 0 | 14 | 0 | 0 | 0 | 0 | 0 | 0 | 0 | 3 | 0 | 1 | 74 | 0 | 0 |
| L_sATL | 0 | 0 | 0 | 0 | 0 | 0 | 0 | 0 | 0 | 0 | 0 | 0 | 3 | 0 | 76 | 0 | 6 | 0 | 0 |
| L_MTG | 0 | 0 | 0 | 0 | 0 | 0 | 0 | 0 | 0 | 0 | 0 | 0 | 0 | 76 | 0 | 11 | 55 | 0 | 0 |
| L_STG | 0 | 0 | 0 | 0 | 0 | 0 | 0 | 0 | 0 | 0 | 0 | 0 | 1 | 0 | 11 | 0 | 17 | 0 | 0 |
| L_SMG | 0 | 0 | 0 | 0 | 0 | 0 | 0 | 0 | 0 | 0 | 0 | 0 | 74 | 6 | 55 | 17 | 0 | 0 | 0 |
| R_IFGpo | 0 | 2 | 0 | 0 | 0 | 2 | 0 | 0 | 0 | 0 | 0 | 0 | 0 | 0 | 0 | 0 | 0 | 0 | 0 |
| R_sATL | 0 | 0 | 0 | 0 | 0 | 0 | 0 | 0 | 0 | 0 | 0 | 0 | 0 | 0 | 0 | 0 | 0 | 0 | 0 |

**Supplementary Figure 16.** Structural connectivity between the nodes of the guilt network at 10% sparsity. The numbers reflect how many individuals were found to have a given connection. Abbreviations: R_mOFC, right medial orbitofrontal cortex; L_ACC, left anterior cingulate cortex; L_SFG, left superior frontal gyrus; L_MeFG_BA6, left medial frontal gyrus (Brodmann area 6); L_MeFG_BA10, left medial frontal gyrus (Brodmann area 10); L_MeFG_BA9, left medial frontal gyrus (Brodmann area 9); L_SCSR, left subgenual cingulate cortex and adjacent septal area; R_SCSR, right subgenual cingulate cortex and adjacent septal area; L_Pcun, left precuneus; L_PCC_POS, left posterior cingulate cortex and parieto-occipital sulcus; L_Cun, left cuneus; R_Ling, right lingual gyrus; L_IFGpo_Ins, left inferior frontal gyrus pars opercularis and insula; L_sATL, left superior anterior temporal lobe; L_MTG, left middle temporal gyrus; L_STG, left superior temporal gyrus; L_SMG, left supramarginal gyrus; R_IFGpo, right inferior frontal gyrus pars opercularis; R_sATL, right superior anterior temporal lobe.

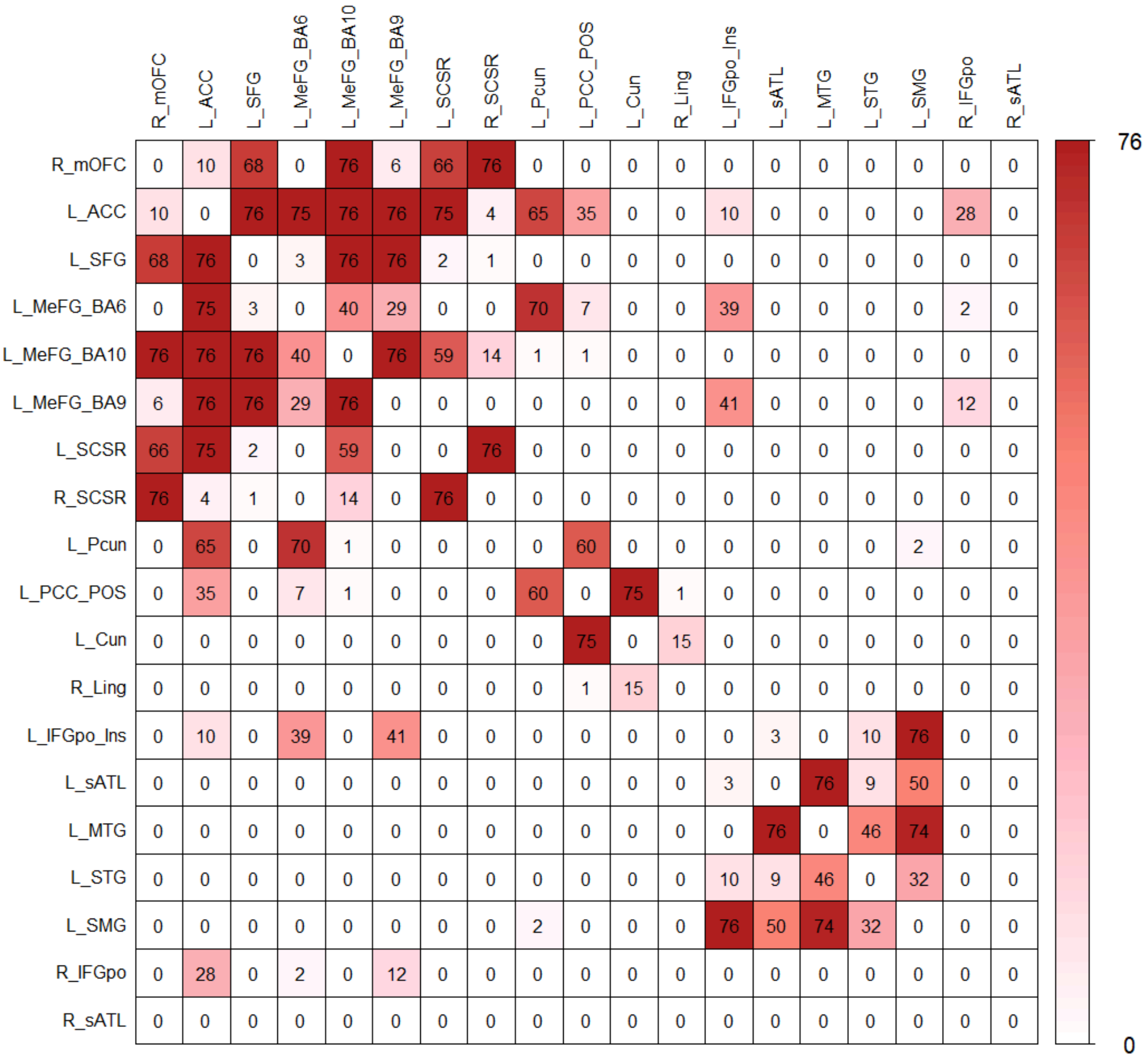


| | R_mOFC | L_ACC | L_SFG | L_MeFG_BA6 | L_MeFG_BA10 | L_MeFG_BA9 | L_SCSR | R_SCSR | L_Pcun | L_PCC_POS | L_Cun | R_Ling | L_IFGpo_Ins | L_sATL | L_MTG | L_STG | L_SMG | R_IFGpo | R_sATL |
|---|---|---|---|---|---|---|---|---|---|---|---|---|---|---|---|---|---|---|---|
| R_mOFC | 0 | 10 | 68 | 0 | 76 | 6 | 66 | 76 | 0 | 0 | 0 | 0 | 0 | 0 | 0 | 0 | 0 | 0 | 0 |
| L_ACC | 10 | 0 | 76 | 75 | 76 | 76 | 75 | 4 | 65 | 35 | 0 | 0 | 10 | 0 | 0 | 0 | 0 | 28 | 0 |
| L_SFG | 68 | 76 | 0 | 3 | 76 | 76 | 2 | 1 | 0 | 0 | 0 | 0 | 0 | 0 | 0 | 0 | 0 | 0 | 0 |
| L_MeFG_BA6 | 0 | 75 | 3 | 0 | 40 | 29 | 0 | 0 | 70 | 7 | 0 | 0 | 39 | 0 | 0 | 0 | 0 | 2 | 0 |
| L_MeFG_BA10 | 76 | 76 | 76 | 40 | 0 | 76 | 59 | 14 | 1 | 1 | 0 | 0 | 0 | 0 | 0 | 0 | 0 | 0 | 0 |
| L_MeFG_BA9 | 6 | 76 | 76 | 29 | 76 | 0 | 0 | 0 | 0 | 0 | 0 | 0 | 41 | 0 | 0 | 0 | 0 | 12 | 0 |
| L_SCSR | 66 | 75 | 2 | 0 | 59 | 0 | 0 | 76 | 0 | 0 | 0 | 0 | 0 | 0 | 0 | 0 | 0 | 0 | 0 |
| R_SCSR | 76 | 4 | 1 | 0 | 14 | 0 | 76 | 0 | 0 | 0 | 0 | 0 | 0 | 0 | 0 | 0 | 0 | 0 | 0 |
| L_Pcun | 0 | 65 | 0 | 70 | 1 | 0 | 0 | 0 | 0 | 60 | 0 | 0 | 0 | 0 | 0 | 0 | 2 | 0 | 0 |
| L_PCC_POS | 0 | 35 | 0 | 7 | 1 | 0 | 0 | 0 | 60 | 0 | 75 | 1 | 0 | 0 | 0 | 0 | 0 | 0 | 0 |
| L_Cun | 0 | 0 | 0 | 0 | 0 | 0 | 0 | 0 | 0 | 75 | 0 | 15 | 0 | 0 | 0 | 0 | 0 | 0 | 0 |
| R_Ling | 0 | 0 | 0 | 0 | 0 | 0 | 0 | 0 | 0 | 1 | 15 | 0 | 0 | 0 | 0 | 0 | 0 | 0 | 0 |
| L_IFGpo_Ins | 0 | 10 | 0 | 39 | 0 | 41 | 0 | 0 | 0 | 0 | 0 | 0 | 0 | 3 | 0 | 10 | 76 | 0 | 0 |
| L_sATL | 0 | 0 | 0 | 0 | 0 | 0 | 0 | 0 | 0 | 0 | 0 | 0 | 3 | 0 | 76 | 9 | 50 | 0 | 0 |
| L_MTG | 0 | 0 | 0 | 0 | 0 | 0 | 0 | 0 | 0 | 0 | 0 | 0 | 0 | 76 | 0 | 46 | 74 | 0 | 0 |
| L_STG | 0 | 0 | 0 | 0 | 0 | 0 | 0 | 0 | 0 | 0 | 0 | 0 | 10 | 9 | 46 | 0 | 32 | 0 | 0 |
| L_SMG | 0 | 0 | 0 | 0 | 0 | 0 | 0 | 0 | 2 | 0 | 0 | 0 | 76 | 50 | 74 | 32 | 0 | 0 | 0 |
| R_IFGpo | 0 | 28 | 0 | 2 | 0 | 12 | 0 | 0 | 0 | 0 | 0 | 0 | 0 | 0 | 0 | 0 | 0 | 0 | 0 |
| R_sATL | 0 | 0 | 0 | 0 | 0 | 0 | 0 | 0 | 0 | 0 | 0 | 0 | 0 | 0 | 0 | 0 | 0 | 0 | 0 |

**Supplementary Figure 17.** Structural connectivity between the nodes of the guilt network at 15% sparsity. The numbers reflect how many individuals were found to have a given connection. Abbreviations: R_mOFC, right medial orbitofrontal cortex; L_ACC, left anterior cingulate cortex; L_SFG, left superior frontal gyrus; L_MeFG_BA6, left medial frontal gyrus (Brodmann area 6); L_MeFG_BA10, left medial frontal gyrus (Brodmann area 10); L_MeFG_BA9, left medial frontal gyrus (Brodmann area 9); L_SCSR, left subgenual cingulate cortex and adjacent septal area; R_SCSR, right subgenual cingulate cortex and adjacent septal area; L_Pcun, left precuneus; L_PCC_POS, left posterior cingulate cortex and parieto-occipital sulcus; L_Cun, left cuneus; R_Ling, right lingual gyrus; L_IFGpo_Ins, left inferior frontal gyrus pars opercularis and insula; L_sATL, left superior anterior temporal lobe; L_MTG, left middle temporal gyrus; L_STG, left superior temporal gyrus; L_SMG, left supramarginal gyrus; R_IFGpo, right inferior frontal gyrus pars opercularis; R_sATL, right superior anterior temporal lobe.
